# Manipulation of plant primary metabolism by leaf-mining larvae in the race against leaf senescence

**DOI:** 10.1101/777334

**Authors:** Mélanie J.A. Body, Jérôme Casas, David Giron

## Abstract

Plants are often considered as suboptimal food for phytophagous insects, requiring them to employ various adaptive mechanisms to overcome food nutritional imbalances. This could include host-plant manipulation and/or symbiotic associations. The extensive reconfiguration of plant primary metabolism upon herbivory, as well as its impact on herbivores, have been largely overlooked, while studies investigating secondary metabolites is extensive. Here, we document how the apple leaf-mining caterpillar *Phyllonorycter blancardella*, a highly-specialized insect which completes development within a restricted area of a single *Malus domestica* leaf over successive different larval feeding modes, maintains nutrient-rich green tissues in its feeding area on green and senescent leaves. For this purpose, we quantified a large number of compounds involved in plant primary metabolism: starch, total soluble sugars, five individual sugars, twenty protein-bound amino acids and twenty free amino acids. Plant alteration can be observed not only on senescing (photosynthetically inactive) but also normal (photosynthetically active) leaf tissues of its host-plant to compensate for detrimental environmental variations. Our results show a differential control of the primary metabolism depending on the larva developmental stage, itself correlated to the fluid-feeding and tissue-feeding modes. Our results also suggest that leaf amino acid alterations favor a faster insect development. Finally, chemical scores indicate that the most growth-limiting essential amino acids are also common to other phytophagous insects and large herbivores, suggesting that these limitations are a general consequence of using plants as food source. We discuss the possible mechanisms responsible for these different manipulative capacities, as well as their ecological implications.

## Introduction

Plants are abundantly present on Earth and are at the basis of most food webs. However, they are often suboptimal food sources for herbivores for three reasons (Mattson, 1980; Schoonhoven et al., 2005; Zhou et al., 2015). First, plant nutrient composition is highly variable according to plant species, organ, season, and other environmental factors (Karasov and Martínez Del Rio, 2007; Schoonhoven et al., 2005; Gündüz and Douglas, 2009). Such variations of plant nutritional quality affect herbivore performance (Karley et al., 2002; Schoonhoven et al., 2005; Larbat et al., 2016). Second, under pathogen infection and herbivore attack, plants mount a defense response and increasing evidence shows an extensive reprogramming of plant metabolism (Schwachtje and Baldwin, 2008; Bolton, 2009; Kerchev et al., 2012). Recent studies demonstrate that hundreds of plant genes involved in defensive pathways are up-regulated during a plant-herbivore interaction and a substantial fraction of them belong to the primary metabolism (Kessler and Baldwin, 2002; Schwachtje and Baldwin, 2008; Kerchev et al., 2012; Zhou et al., 2015). Third, nitrogen is considered to be the most limiting element for the growth of most herbivores (Mattson, 1980; Van Zyl and Ferreira, 2003; Barbehenn et al., 2013; Zhou et al., 2015) and there is ample evidence that amino acid acquisition (including protein-bound and free amino acids) does not match the proportion of essential amino acids (EAAs) they require. Indeed, nitrogen concentration in plants is generally low compared with animal tissues (Bernays and Chapman, 1994; Karasov and Martínez Del Rio, 2007). Thus, the challenge for herbivore is to find and eat foods that contain the best blend of nutrients (specific composition in specific tissues) and at the most adequate concentration (overall plant sugar and/or amino acid content) (Chown and Nicolson, 2004; Schoonhoven et al., 2005; Klowden 2007). Otherwise, acquisition of unbalanced food could impair key physiological processes such as protein synthesis, and/or would require sophisticated and costly adaptations (Horie and Watanabe, 1983; Karowe and Martin, 1989; Douglas et al., 2001; Suzuki et al., 2009).

Because of the major nutritional discrepancy which clearly exists between plant nutrient supply and herbivore nutritional requirements, some species have developed different strategies such as symbioses with microorganisms that supplement insects with limiting EAAs (Douglas, 2006, 2009, 2013, and references therein). Alternatively, many folivorous organisms are selective feeders, consuming certain tissues or cell types and rejecting others. This gives them the unique possibility to consume high-quality tissues in an otherwise low-quality plant or plant organ. Many leaf-mining insects, for example, consume internal mesophyll cells and do not eat epidermis and/or vascular tissues (Hering, 1951; Kimmerer and Potter, 1987; Scheirs et al., 2001; Body et al., 2015). Other insects, such as aphids, are in more privileged position than chewing insects, because nearly all nitrogen-containing compounds in phloem sap can be utilized (Schoonhoven et al., 2005); still, they often, if not always, rely on symbionts to provide them with key EAAs (Douglas et al., 2001; Wu et al., 2006). Finally, other organisms modify the nutritional quality of the host-plant for their own needs by creating specialized tissues to feed on (Hartley, 1998; Nakamura et al., 2003; Koyama et al., 2004; Dardeau et al., 2015), by upregulating protein and/or sugar synthesis *in situ* (Saltzmann et al., 2008), and/or by modifying source-sink relationships leading to nutrient translocation towards the insect’s feeding site (Larson and Whitham, 1991). The flow of nutrients towards the infection site, which thus behaves as a metabolic sink, has obvious advantages to insects by fueling energetic requirements for growth, survival and reproduction (Stone and Schönrogge, 2003; Giron et al., 2007, 2016; Body et al., 2013). This has been clearly demonstrated, for example, with radioactively labelled nutrients that are preferentially transported and accumulated in specific areas exploited by herbivorous insects (Larson and Whitham, 1991). For example, some gall-inducing insects enhance their nutritional environment by increasing the abundance of plant free amino acids and/or sugars through alteration of their synthesis and/or transport (Larson and Whitham, 1991; Eleftherianos et al., 2006; Saltzmann et al., 2008; Dardeau et al., 2015). Interestingly, improved nutrition does not necessarily imply increased nutrient concentrations, but an adequate composition and/or quantity, leading to a finely tuned regulation of nutrient contents in some biological systems, such as for gall-inducing insects *Neuroterus quercus-baccarum* and *Andricus lignicola* (Hartley and Lawton, 1992; Diamond et al., 2008).

Although qualitative and quantitative variation in *primary* plant compounds, as opposed to the well know secondary compounds, can have profound effects on phytophagous insect preferences and performances, little is known about plant *primary* metabolism reconfiguration after herbivore attack (Machado et al., 2013, 2015) and data concerning specific feeding-guild with high nutritional constraints are scarce (Schoonhoven et al., 2005). Plant nutrient availability is particularly crucial for endophagous insects such as stem-borers, gall-inducers, and leaf-miners. Their entire life-cycle is indeed usually restricted to the same plant organ without diet switching capacities as observed for free-living external-feeding insects (Hering, 1951; Stone and Schönrogge, 2003; Body et al., 2015). As such, they are the most appropriate feeding-guild in which to examine nutritional constraints imposed by the plant and adaptive strategies adopted by the insect to overcome these constraints. Additionally, changes in the ability and/or need to manipulate the host-plant physiology is expected to vary across insect feeding-mode and developmental stages (Schoonhoven et al., 2005), as herbivorous insects possess a diverse range of feeding habits which differ in the nutritive source they have access to and as nutritional requirements change during the insect development. This prompted us to carry out a detailed biochemical investigation of plant carbohydrate and amino acid profiles in the host-plant *Malus domestica* following attack by the leaf-mining caterpillar *Phyllonorycter blancardella* according to the strongly differing larval feeding-modes.

*P. blancardella* leaf-mining moth presents an important degree of specialization for its host-plant and a complete development within a restricted area of a single leaf (Hering, 1951; Pottinger and LeRoux, 1971; Connor and Taverner, 1997; Body et al., 2015). In this species, first instars are ‘fluid-feeders’ while the two last instars are ‘tissue-feeders’ due to a hypermetamorphosis of their mouthparts (characteristic changes in morphology and habit between two successive instars) (Body et al., 2015). Previous studies on this biological system focused on later tissue-feeding instars only (Giron et al., 2007; Kaiser et al., 2010; Body et al., 2013; Zhang et al., 2016, 2017). This leaf-miner is able to induce an accumulation of cytokinins in mined tissues which is responsible for the preservation of photosynthetically active green tissues (‘green-island’ phenotype) at a time when leaves are otherwise turning yellow (i.e. senescent leaves) (Engelbrecht et al., 1969; Giron et al., 2007; Kaiser et al., 2010; Body et al., 2013; Zhang et al., 2017). Host-plant physiological alterations occur through an unexpected association with endosymbiotic bacteria (*Wolbachia*) (Kaiser et al, 2010; Body et al., 2013; Giron and Glevarec, 2014; Gutzwiller et al., 2015). The correlation between the green-island phenotype and *Wolbachia* infections has also been highlighted in numerous species of Gracillariidae leaf-mining moths (Gutzwiller et al., 2015). The strong reprogramming of the plant phytohormonal balance (CKs, JA, SA, ABA; Body et al., 2013; Zhang et al., 2016) reported is associated with the regulation of sugar content (Body et al., 2013), inhibition of leaf senescence (Kaiser et al., 2010) and mitigation of plant direct and indirect defense (Giron et al., 2016; Zhang et al., 2016).

To gain insight into the extend of nutrient regulation and subsequent consequences for *P. blancardella* leaf-miners developing under harsh environmental conditions, we first characterized how larvae with different feeding-modes (fluid- vs. tissue-feeding instars) impact starch, soluble sugar, and protein-bound and free amino acid profiles at their feeding site both on green and on yellow senescing leaves. Senescence is a particularly decisive moment for these insects due to the profound alteration of the plant physiology including the nutrient (sugars and degraded proteins) remobilization to roots. In the Fall, when nutrients become too low, the insect growth usually stops (Edwards and Wratten, 1980). If *P. blancardella* fluid-feeding larvae fail to reach the tissue-feeding stage before low Fall temperatures, caterpillars will not be able to complete their development and to pupate which lead to an increased mortality rate (Pottinger and LeRoux, 1971).

Sugars and amino acids play a crucial role in life maintenance as a large source of energy for insects, as a structural component of cuticular chitin, and as feeding and oviposition stimulants (Dadd, 1985; Nation, 2002). Estimating if observed plant physiological modifications are beneficial for the insect requires us to measure the impact of various food compositions on insect fitness-related traits. The lack of artificial diet for bioassays with leaf-mining insects impairs such analyses. However, the use of indirect method such as “chemical scores” allows to determine growth-limiting EAAs for herbivores (Van Zyl and Ferreira, 2003; Barbehenn et al., 2013). We thus estimated dietary EAA requirements of the leaf-mining larvae by determining the whole-body EAA composition of the insect and confronting these results with plant chemical profiles (Rock, 1972; Van Zyl and Ferreira, 2003; Anderson et al., 2004; Barbehenn et al., 2013). In this study, we expected (*i*) a finely-tuned nutrient accumulation (sugars and amino acids) at the feeding site to meet insect nutritional requirements, and (*ii*) a differential control of the nutritional content depending on the larva developmental stage as the impact of herbivory on plant tissues is correlated to leaf-miner feeding-mode.

## Materials and Methods

### Study species and sampling sites

The experiments were conducted on *Malus domestica* (Borkh. 1803) (Rosaceae) apple-tree leaves naturally infected by the spotted tentiform leaf-miner, *Phyllonorycter blancardella* (Fabricius, 1781) (Lepidoptera: Gracillariidae). This leaf-miner species is a polyvoltine microlepidopteran widely distributed in Europe whose first three instars are fluid-feeders and two following instars are selective tissue-feeders (Pottinger and LeRoux, 1971; Body et al., 2015).

Both green and yellow mined leaves (only one mine per leaf), and unmined green and yellow leaves (an adjacent neighboring leaf), were simultaneously collected in the field between 08:00 a.m. and 10:00 a.m. in early Falls 2009-2011 on 16-18-year-old apple-trees (“Elstar” varieties), in a biologically managed orchard in Thilouze, France (47° 14’ 35” North, 0° 34’ 43” East). Collected leaves and associated larvae were immediately kept and dissected on ice, and stored at −80 °C until analysis. The synchronization of sampling is crucial as levels of sugars, for example, greatly vary during the day and among trees. This requires collecting green and yellow leaves (mined leaves and their unmined control leaves) simultaneously and on the same tree to make sure that physiological differences observed are due to the impact of the leaf-miner on plant tissues and not to phenological changes in the tree.

In order to study spatial (mined vs. unmined areas) and temporal (senescence) variations of starch, total and individual sugar, and protein-bound and free amino acid concentrations, mined tissues (zone M; leaf-mining insects and faeces were removed from the mine), ipsilateral tissues (zone UM^1^; leaf tissues on the same side of the main vein as the mine), and contralateral tissues (zone UM^2^; leaf tissues on the opposite side of the main vein as the mine) were dissected both on green and on yellow leaves (Figure 1.1). Non-infected green and yellow leaves (zone UM^3^) were also dissected as previously and used as a control (Giron et al., 2007; Body et al., 2013).

**Figure 1.**
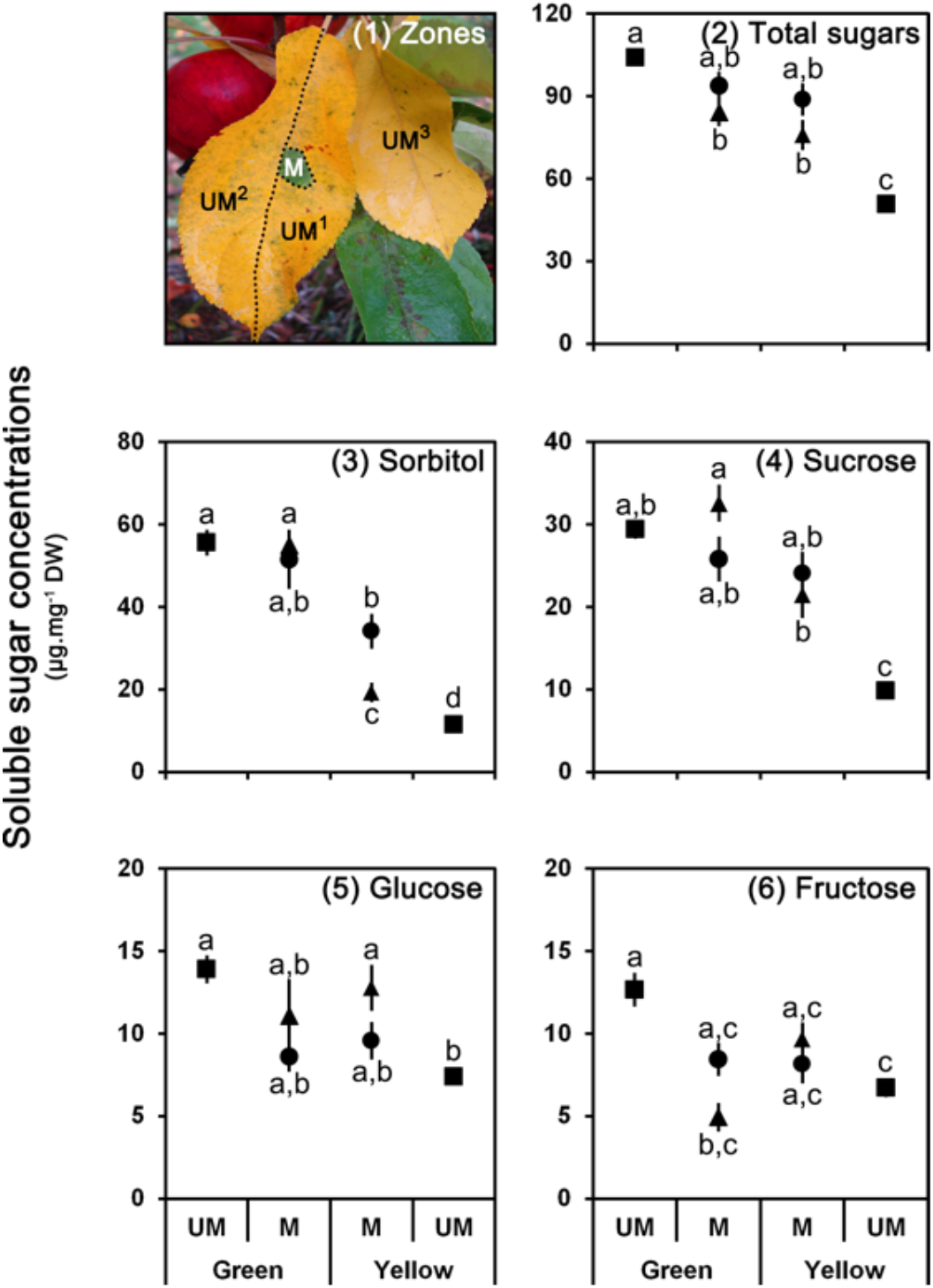
Amounts of sugars in leaf tissues. (**1**) Picture of a yellow mined leaf: mined tissues (M zone), ipsilateral unmined tissues (UM^1^ zone), contralateral unmined tissues (UM^2^ zone), and unmined control leaf (UM^3^ zone). Amounts of (**2**) total soluble sugars, (**3**) sorbitol, (**4**) sucrose, (**5**) glucose and (**6**) fructose in unmined tissues (UM; squares) and mined tissues (M) for fluid-feeding (L3 instar; triangles) and tissue-feeding (L5 instar; circles) larvae both on green and on yellow leaves. Data are expressed as μg per mg of leaf dry weight (DW) and presented as average ± S.E.M. Statistical differences between averages are shown by different letters (a, b, c, d) (see Supplement 2 for details).

### Sample preparation

Each leaf sample was lyophilized (Bioblock Scientific Alpha 1-4 LD plus lyophilizator) according to the following cycle: primary desiccation of 1 hour at −10°C and 25 mbar, and a secondary dessiccation overnight at −76°C and 0.0010 mbar. Samples were then ground(ed?) in liquid nitrogen after in order to have an extra-fine leaf powder. Similar amounts (5 mg) of mined (M), ipsilateral (UM^1^), contralateral (UM^2^) and control (UM^3^) plant tissues were used to allow qualitative and quantitative comparisons (Sartorius micro-balance model 1801-001, Sartorius SA, Palaiseau, France).

### Carbohydrate quantification

Leaf samples used for this experiment were as follow: *green leaves*: N = 15 for fluid-feeding instars, N = 25 for tissue-feeding instars, N = 120 for unmined green control; *yellow leaves*: N = 15 for fluid-feeding instars, N = 25 for tissue-feeding instars, N = 120 for unmined yellow control.

#### Carbohydrate extraction

Prior to colorimetric quantifications, chlorophyll and other pigments were removed from leaf tissues (5 mg) with acetone (100 %) until complete elimination of natural coloration. Sugars were then extracted with vortex agitation for 30 sec at room temperature in 1 mL aqueous methanol (80 %) (Fisher Scientific; Hampton, New Hampshire, USA). After centrifugation at 1500 rpm, soluble sugars remained dissolved in the supernatant and were used for: (*i*) total soluble sugars quantification by colorimetric assays following Van Handel’s protocol (1985) as modified by Giron et al. (2002) for microsamples, and (*ii*) for characterization of individual sugars by capillary electrophoresis using a modification of Rovio’s protocol optimizing sugars separation and data reproducibility (Rovio et al., 2007; see Body et al., 2013 for details). Starches remained in the pellet were quantified using a modification of colorimetric techniques developed by Hansen and Møller (1975), Marshall (1986) and Oren et al. (1988). The anthrone reagent consisted of 1.0 g of anthrone (Sigma Aldrich; St. Louis, Missouri, USA) dissolved in 500 mL of concentrated sulfuric acid (Fisher Scientific; Hampton, New Hampshire, USA) added to 200 mL of MilliQ water (Merck Millipore; Billerica, Massachusetts, USA).

#### Starch quantification

Leaf material (pellet) kept for starch quantification was suspended in 1 mL of hydrochloric acid 1.1 % (Fisher Scientific; Hampton, New Hampshire, USA), vortexed and placed in a water bath at 90 °C for 30 min to extract and hydrolyze starch into glucose molecules. In a new set of tubes, 1 mL of anthrone reagent was then added to 35 μL of the extraction solution for each sample. The tubes were reheated at 90 °C for 15 min, cooled down at 0 °C for 5 min, vortexed. Absorbance was then read in a spectrophotometer at 630 nm (DU®-64 spectrophotometer; Beckman, Villepinte, France). For starches, calibration curves that allowed us to transform absorbance into concentrations were made with standard glucose (Sigma Aldrich; St. Louis, Missouri, USA).

#### Total soluble sugar quantification

For each sample, 100 μL of initial aqueous methanol supernatant were transferred into a borosilicate tube (16 × 100 mm; Fisher Scientific; Hampton, New Hampshire, USA) and placed in a water bath at 90 °C to evaporate the solvent down to a few microlitres. After adding 1 mL of anthrone reagent, the tubes were placed in a water bath at 90 °C for 15 min, cooled down at 0 °C for 5 min, vortexed and then read in a spectrophotometer at 630 nm (DU®-64 spectrophotometer; Beckman, Villepinte, France). For total soluble sugars, calibration curves were corrected for the underestimation of sugar alcohols using a sugar mixture (sorbitol, trehalose, sucrose, glucose and fructose; Sigma Aldrich; St. Louis, Missouri, USA) (Body et al., 2013; Body et al., 2018) close to the composition of mined and unmined tissues both on green and on yellow leaves (Figure 2).

**Figure 2.**
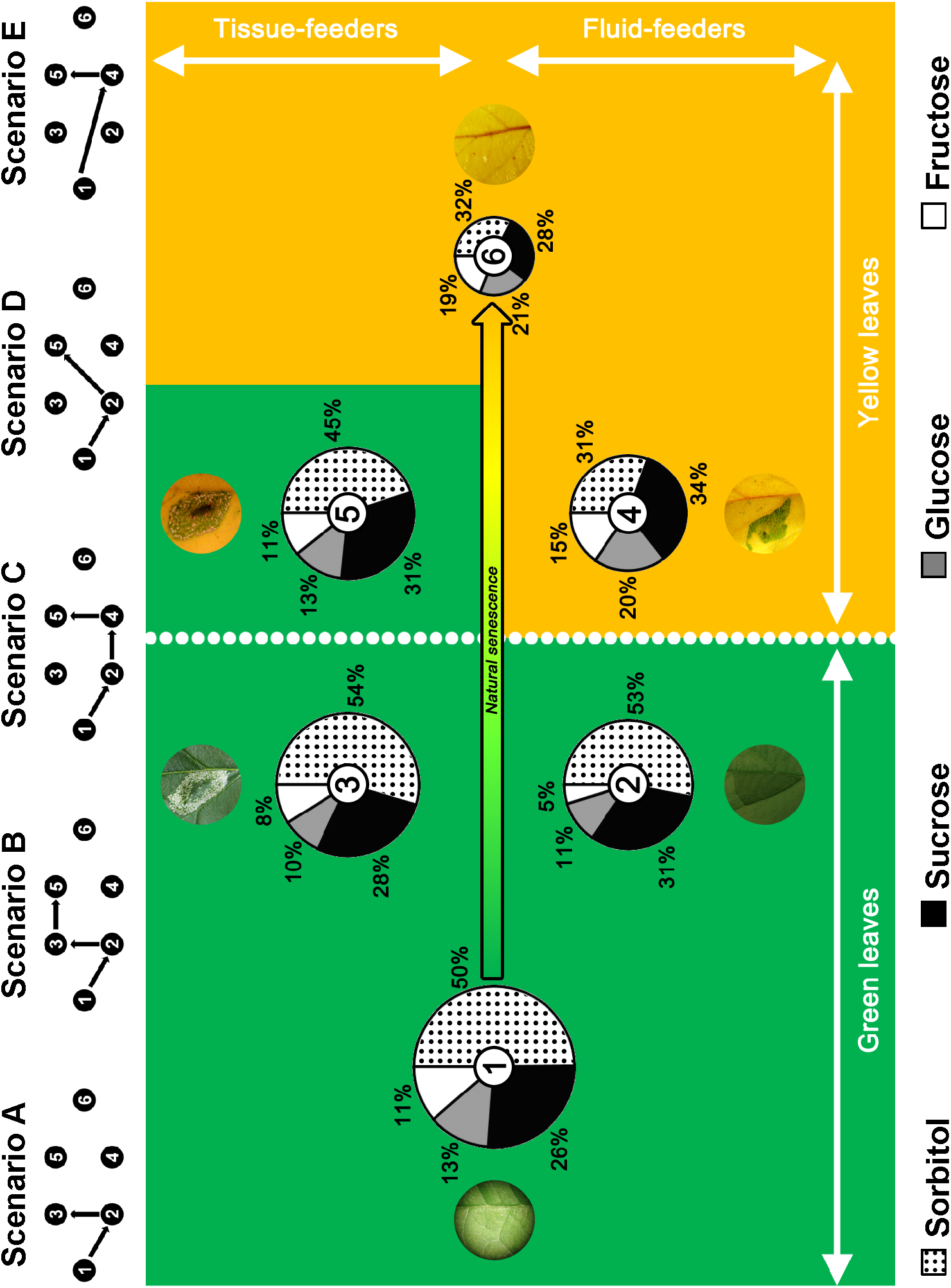
Sugar composition of leaf tissues according to the leaf-mining process. Sugar composition of unmined (**1** and **6**) and mined tissues for fluid-feeding (**2** and **4**) and tissue-feeding (**3** and **5**) larvae both on green (**1** to **3**) and on yellow (**4** to **6**) leaves. Size of pies represents the total amount of sugars. The green background symbolizes similarities of sugar quantities and compositions with an unmined green leaf. The yellow background symbolizes similarities of sugar quantities and compositions with an unmined yellow leaf. The dynamic of the leaf-mining process is presented through five possible scenarii, eggs being always laid on green leaves and yellowing of leaves occurring at different possible larval development stages. Status 1: Unmined green leaf. Status 2: Fluid-feeding larva on green leaf. Status 3: Tissue-feeding larva on a green leaf. Status 4: Fluid-feeding larva on a yellow leaf. Status 5: Tissue-feeding larva on a yellow leaf. Status 6: Unmined yellow control. See Supplement 2 for statistical analysis.

#### Individual sugars quantification and capillary electrophoresis conditions

Remaining supernatants from previous extractions were used for individual sugars assay by capillary electrophoresis (Proteome lab™ PA800, 32Karat data acquisition software; Beckman Coulter; Brea, California, USA) (Rovio et al., 2007). For each sample, 500 μL of supernatant were filtered at 0.45 μm (Whatman®; Maidstone, United Kingdom) and then lyophilized (Bioblock Scientific Alpha 1-4 LD plus lyophilizator). Right before analysis, sugars were resuspended by vortex agitation in 18 μL of MilliQ water (Merck Millipore; Billerica, Massachusetts, USA) and 2 μL melezitose 50 mM (Sigma Aldrich; St. Louis, Missouri, USA), as an internal standard, and then centrifuged briefly. For sugar separation, we used a raw silica capillary of 50 μm internal diameter and 67 cm total length (Beckman Coulter; Brea, California, USA). Samples (2.77 nL) were injected and maintained at 15 °C during all the separation which occurred at 16 kV (typical current 90 μA) and lasted for maximum 60 min. The electrolyte was composed of NaOH 130 mM and Na_2_HPO_4_·12H_2_O 36 mM, pH 12.6 (Rovio et al., 2007). The UV detection wavelength was 264 nm. The control solution was composed of 7 standard sugars at 5 mM: D-(+)-xylitol, D-(+)-sorbitol, D-(+)-trehalose, D-(+)-melezitose (internal standard), D-(+)-sucrose, le D-(+)-glucose and D-(+)-fructose. All sugar standards were purchased at Sigma Aldrich (St. Louis, Missouri, USA). Data were processed with the 32Karat™ Software (version 7.0) which allowed for the determination of retention times and peak areas for each reference sugars (standards sugars).

#### Data analysis

Individual sugar analysis revealed the presence of only five main sugars: sorbitol (retention time: 17.5 min), trehalose (RT: 17.7 min), sucrose (RT: 21.0 min), glucose (RT: 27.8 min), and fructose (RT: 29.3 min) [internal standard, melezitose, RT: 20.3 min]. However, as trehalose had a low concentration, was partially co-elued with another sugar (sorbitol), and highly variable, data were not included in the present study in order to avoid large estimation errors for this sugar. Unlike trehalose, sorbitol was highly concentrated in apple-tree leaves leading to an accurate/reliable quantification with a negligible estimation error due to this partial peak co-elution.

In order to compare the biochemical composition of mined and unmined tissues, we had to take into account the withdrawal of sugar-rich mesophyll tissues by leaf-mining insects and the over-representation of sugar-free epidermis in the mined tissue samples. For this purpose, gravimetry was used to estimate the amount of mesophyll eaten by the larva and to correct biochemical data of leaf tissues accordingly (see Supplement 1 for details). All data presented are thus corrected for the amount of tissues eaten by larvae as this parameter was highly significant for all sugars (*Student’s paired t-test*: *P* < 0.001 for total sugars, sucrose, glucose and fructose; *Wilcoxon paired test*: *P* < 0.01 for sorbitol).

### Amino acids quantification

Two sets of leaf samples were used for this experiment (one for protein-bound amino acid and one for free amino acid) and were as follow for each set: *green leaves:* N = 15 for fluid-feeding instars, N = 30 for tissue-feeding instars, N = 135 for unmined green control; *yellow leaves:* N = 15 for fluid-feeding instars, N = 30 for tissue-feeding instars, N = 135 for unmined yellow control.

#### GC-MS conditions

Standard physiological amino acids at a concentration of 200 nmol.ml^−1^, norvaline used as internal standard and reagents were supplied in the EZ:faast GC-MS kit for amino acid analysis (Phenomenex; Torrance, California, USA). A Perkin Elmer AutoSystemXL gas chromatograph was directly coupled to a Perkin Elmer TurboMass mass spectrometer (Perkin Elmer; Waltham, Massachusetts, USA). A 10 m × 0.25 mm ZB-AAA column from EZ:faast kit of Phenomenex (Torrance, California, USA) was used. The carrier gas helium (Air Liquide; Paris, France) flow was kept constant at 1.1 mL/min. The oven temperature program was a 30 °C/min ramp from 110 °C to 320 °C. The temperature of the injection port was 250 °C. The MS temperatures were as follows: ion source (electronic impact) 200 °C, and inlet line temperature 310 °C. The scan range was 3.5 scans/s and mass detected between 45 and 450. Under these conditions a 2 μL sample was injected in splitless mode during 30 sec.

#### Free amino acid extraction

On a subset of samples, free amino acids from leaf or insect samples were extracted with vortex agitation for 30 sec at room temperature in 1 mL acetonitrile 25 % (Fisher Scientific; Hampton, New Hampshire, USA) in hydrochloric acid 0.01 N (1:3, v:v) (Fisher Scientific; Hampton, New Hampshire, USA).

#### Protein-bound amino acid extraction

On another subset of samples, proteins were hydrolyzed into their protein-bound amino acids in a sealed glass tube at 150 °C for 2 h with 500 μL methanesulfonic acid 4 M (Fisher Scientific; Hampton, New Hampshire, USA) after flushing out air with a gentle stream of nitrogen gas. Unlike HCl hydrolysis, methanesulfonic acid hydrolysis allowed the determination of all residues, including tryptophan (Chiou and Wang, 1988; Fountoulakis and Lahm, 1998). Following hydrolysis, the hydrolysates were partially neutralized with 1 mL sodium carbonate 1 M. Prior analysis, samples were transferred in a 1.5 mL Eppendorf tube and pH were checked to be in the range 1.5-5.0.

#### Amino acid derivatization and quantification

One hundred μL of each sample (protein-bound or free amino acid extract) were pipetted into a glass vial and 100 μL of internal standard (norvaline at 200 nmol.mL^−1^) were added. After solid-phase extraction (SPE), sample derivatization was performed as described in Phenomenex kit protocol. Samples were then concentrated under a gentle stream of nitrogen gas (Air Liquide; Paris, France) under 5 μL and immediately injected into the GC-MS system.

#### Data analysis

Chromatograms were analyzed with the TurboMass software (version 5.4.2; Perkin Elmer; Waltham, Massachusetts, USA). Retention times and characteristic ions were as follow: alanine (RT: 1.5 min; ions: 130, 70), glycine (RT: 1.6 min; ions: 116, 74), valine (RT: 1.8 min; ions: 158, 72), leucine (RT: 2.0 min; ions: 172, 86), isoleucine (RT: 2.1 min; ions: 172, 130), threonine (RT: 2.3 min; ions: 160, 101), serine (RT: 2.3 min; ions: 203, 146, 101), proline (RT: 2.4 min; ions: 243, 156), asparagine (RT: 2.5 min; ions: 155, 69), arginine (RT: 3.1 min; ion: 303), aspartate (RT: 3.1 min; ions: 216, 130), methionine (RT: 3.2 min; ions: 277, 203), glutamate (RT: 3.5 min; ions: 230, 170, 84), phenylalanine (RT: 3.5 min; ions: 206, 190, 148), cysteine (RT: 3.9 min; ions: 248, 206, 162), glutamine (RT: 4.2 min; ions: 187, 84), lysine (RT: 4.8 min; ions: 170, 128), histidine (RT: 5.0 min; ions: 282, 168), tyrosine (RT: 5.3 min; ions: 206, 107), tryptophan (RT: 5.6 min; ion: 130), and cystine (RT: 6.4 min; ions: 248, 216) [internal standard, norvaline, RT: 1.9 min; ion: 158]. The elution did not allow the separation of all amino acids (threonine was co-eluted with serine, arginine with aspartate, and glutamate with phenylalanine), which prevent from determining accurately their limitation in mined tissues for leaf-mining larvae. The co-eluted peaks are, unfortunately, undistinguishable. In order to determine changes in leaves in essential (EAA) and non-essential (NEAA) amino acids separately, concentrations of co-eluted EAA and NEAA were thus estimated, for simplicity, by dividing co-eluted peaks by two, assuming that both amino acids were present in the same quantities in each sample.

### Limiting essential amino acids in mined tissues

The plant nutritional quality represented by the amino acid pool available can be estimated based on which EAAs [valine (Val), leucine (Leu), isoleucine (Ile), threonine (Thr), arginine (Arg), methionine (Met), phenylalanine (Phe), lysine (Lys), histidine (His), and tryptophan (Trp), that animals cannot synthesize *de novo*] have the lowest abundance relative to the composition required by an herbivore (growth-limiting EAAs) (Van Zyl and Ferreira, 2003; Barbehenn et al., 2013). All calculations were performed using total amino acid content which is composed of protein-bound and free amino acids. The composition of each EAA was calculated as (μg EAA in sample ÷ μg total amino acids in sample) × 100. Total amino acids are composed of protein-bound and free amino acids. The chemical score for each EAA was calculated as the composition of the EAA in mined tissues ÷ composition of the EAA in the leaf-miner (Barbehenn et al., 2013). Chemical scores were calculated using the mean values from mined tissues and whole leaf-mining insect. For example, the chemical score of histidine for tissue-feeding instars on yellow leaves is calculated as follows: His in the leaf mined tissues (0.59 % of total amino acids) ÷ His in the body of the leaf-mining insect (62.32 % of total amino acids) = 0.01. The EAAs with chemical scores lower than 1 were defined as limiting EAAs (Barbehenn et al., 2013).

### Statistical analysis

Statistical analyses were performed using R version 3.2.1 and RStudio version 0.99.467 (The R Foundation for Statistical Computing, Vienna, Austria). Preliminary statistical analysis showed that the nutrient content of unmined zones (UM^1^, UM^2^, UM^3^) were identical (*Behrens-Fisher test*: *P* > 0.05), allowing for the statistical analysis of mined (M) versus unmined tissues (UM; ipsilateral + contralateral + non-infected leaf) in the result section. The total soluble sugar, individual sugar, protein-bound and free amino acid contents were analyzed separately using *Kruskal-Wallis test* and *Behrens-Fisher post-hoc test*. The sugar composition from different subsets of leaves/tissues was analyzed using multivariate analysis of variance (*MANOVA*). For *MANOVA* analyses, we used the *Pillai’s test statistic*. Where significant effects were observed, post-hoc comparisons were performed. All nutrient quantities are presented in μg per mg of dry weight (DW) as average ± standard error of the mean (S.E.M.).

### Abbreviations

*EAAs* essential amino acids, *NEAAs* non-essential amino acids, *Ala* alanine, *Arg* arginine, *Asn* asparagine, *Asp* aspartate, *Cys* cysteine, *Gln* glutamine, *Glu* glutamate, *Gly* glycine, *His* histidine, *Ile* isoleucine, *Leu* leucine, *Lys* lysine, *Met* methionine, *Phe* phenylalanine, *Pro* proline, *Ser* serine, *Thr* threonine, *Trp* tryptophan, *Tyr* tyrosine, *Val* valine.

## Results

### Alteration of primary metabolism in mined tissues

To evaluate the impact of leaf-mining larvae on the nutritional value of plant tissues, we conducted experiments investigating carbon and nitrogen contents (starch, total soluble and individual sugars, and protein-bound and free amino acids) of mined tissues compared to unmined control tissues. See Supplement 2 for a detailed statistical analysis concerning carbohydrates and Supplements 3 and 4 concerning amino acids. The data are presented in Figures 1 and 2 for carbohydrates and in Figures 3, 4 and 5 for amino acids.

**Figure 3.**
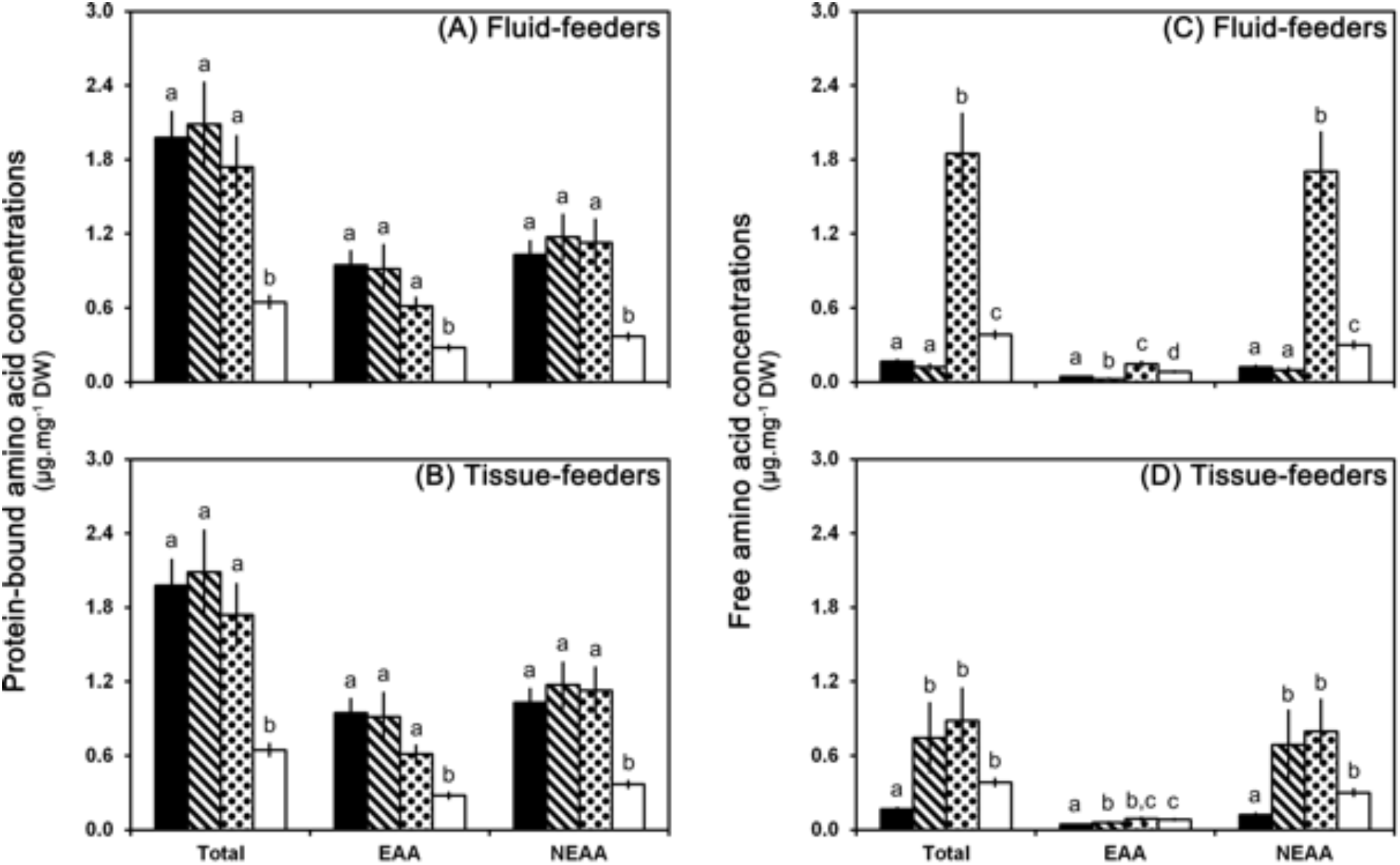
Amino acid composition (total, essential and non-essential amino acids) of mined and unmined tissues both on green and on yellow leaves. Upper panels (**A** and **C**) show data obtained for fluid-feeding instars of the leaf-mining insect *Phyllonorycter blancardella*. Lower panels (**B** and **D**) show data obtained for tissue-feeding instars. Left panels (**A** and **B**) show data for protein-bound amino acids. Right panels (**C** and **D**) show data for free amino acids. Mined tissues on green leaves are represented by the stripe pattern and on yellow leaves by the dot pattern. Unmined green controls are in black and unmined yellow controls are in white. Data are presented as average ± S.E.M. and expressed in μg per mg of leaf dry weight. Statistical differences between averages are shown by different letters (a, b, c, d) (see Supplements 3 and 4 for statistical analysis).

**Figure 4.**
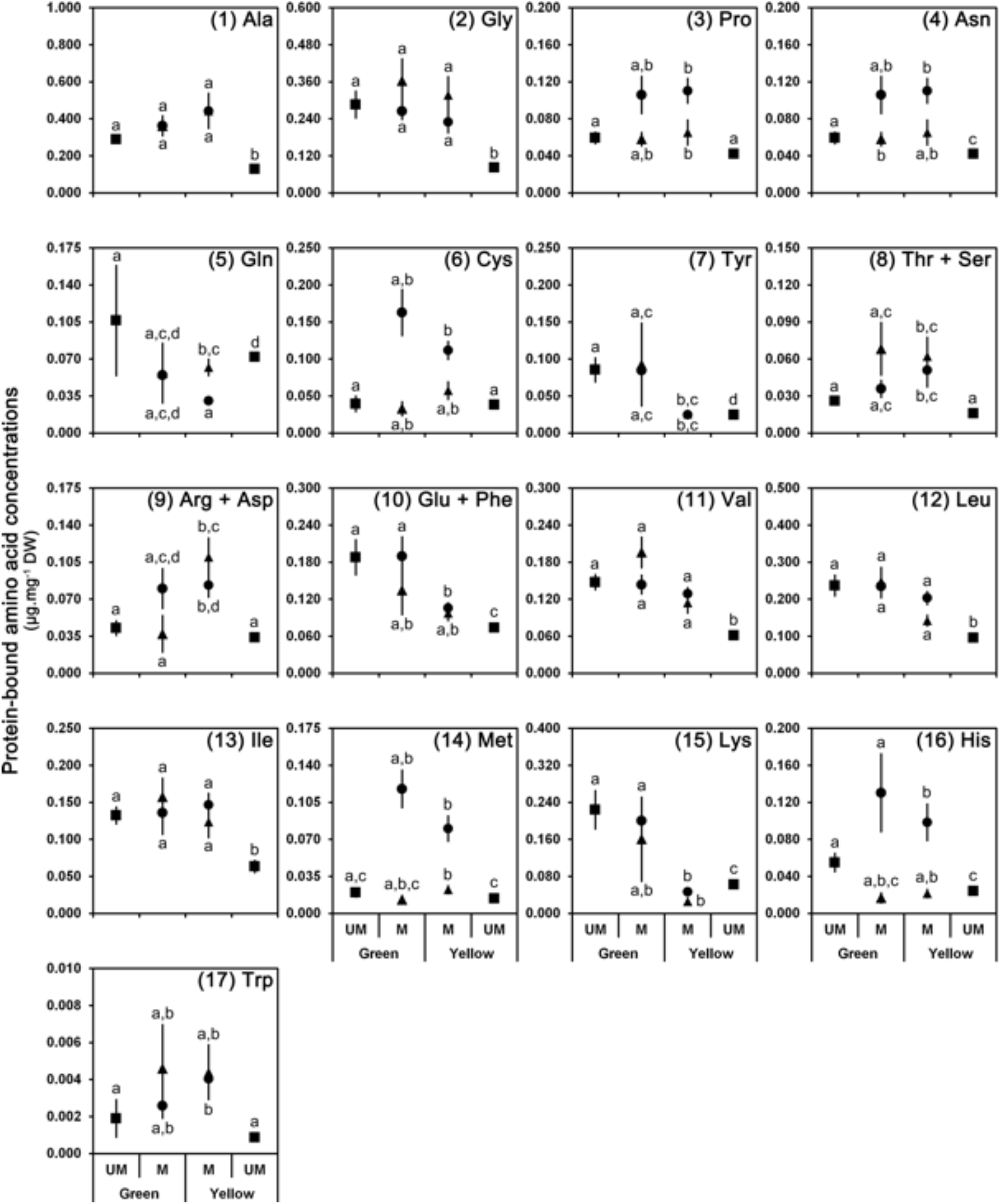
Protein-bound amino acid profiles in mined (M) and unmined (UM; squares) tissues both on green and on yellow leaves infected by the leaf-mining insect *Phyllonorycter blancardella* (fluid-feeding (triangles) vs. tissue-feeding (circles) instars). Panels **1**-**7** are NEAA, panels **8**-**10** are co-eluted NEAA and EAA, panels **11**-**17** are EAA. Data are presented as average ± S.E.M. and expressed in μg per mg of leaf dry weight. Statistical differences between averages are shown by different letters (a, b, c, d) (see Supplement 3 for statistical analysis).

**Figure 5.**
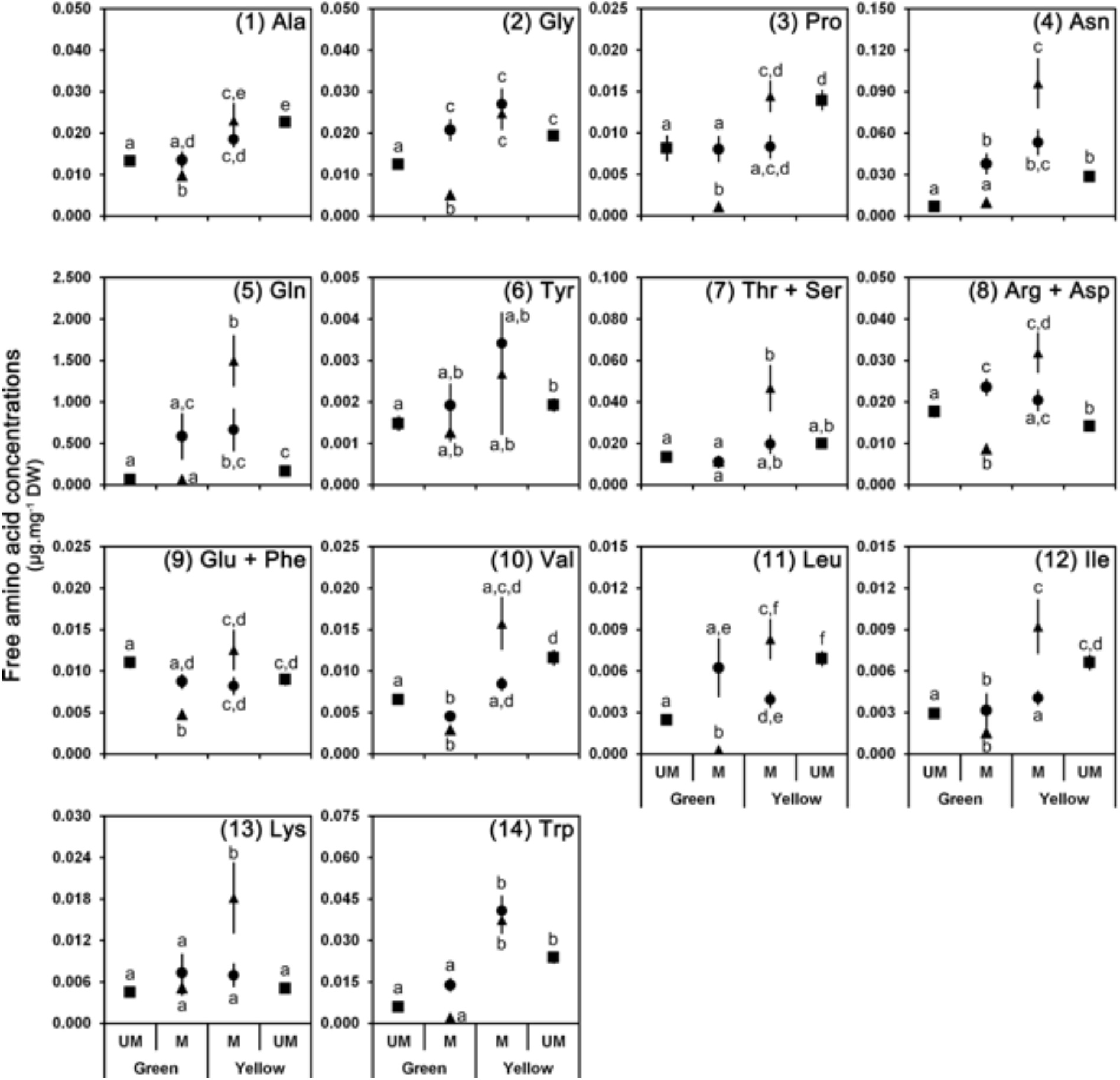
Free amino acid profiles in mined (M) and unmined (UM; squares) tissues both on green and on yellow leaves infected by the leaf-mining insect *Phyllonorycter blancardella* (fluid-feeding (triangles) vs. tissue-feeding (circles) instars). Panels **1**-**6** are NEAA, panels **7**-**9** are co-eluted NEAA and EAA, panels **10**-**14** are EAA. Data are presented as average ± S.E.M. and expressed in μg per mg of leaf dry weight. Statistical differences between averages are shown by different letters (a, b, c, d, e, f) (see Supplement 4 for statistical analysis).

#### Sugars

Individual sugar analysis revealed the presence of five main sugars: sorbitol, trehalose, sucrose, glucose, and fructose. Trehalose data were not included (see Material and Methods section). Senescence leads to a drastic reduction of the total amount of sugars (*green control*: 104.0 ± 2.4 μg.mg^−1^ DW; *yellow control*: 50.6 ± 1.3 μg.mg^−1^ DW; *Behrens-Fisher test*: *P* < 0.001) and an alteration of the specific sugar composition (*MANOVA*: *F*_4,75_ = 18.49, *P* < 0.001; Figures 1 and 2 – green controls vs. yellow controls; Supplement 2A, left column). Amounts of total and individual sugars in tissues mined by fluid-feeders are similar but not statistically identical on green and on yellow leaves (Supplement 2A, middle column), whereas tissues mined by tissue-feeders remain statistically similar during senescence (Supplement 2A, right column).

Composition analysis (Figure 2) showed that, on green leaves, tissues mined by fluid-(Supplements 2B and 2D, left column) and tissue-feeders (Supplements 2C and 2D, left column) are similar to unmined green controls for both total sugar content and specific sugar composition. By contrast, on yellow leaves, the sugar composition of tissues mined by fluid-feeders is identical to the composition of unmined yellow controls (Supplement 2B, right column), whereas for tissues-feeders, the sugar composition of mines is similar to unmined green controls (Supplement 2C, right column).

#### Starch

Senescence leads to the decrease of starch content from 73.54 ± 3.00 μg.mg^−1^ DW in green controls to 53.28 ± 2.27 μg.mg^−1^ DW in yellow controls (Supplement 2A, left column). Starch amounts in leaf tissues mined by fluid-feeders (*green leaves*: 34.57 ± 4.25 μg.mg^−1^ DW; *yellow leaves*: 40.42 ± 3.96 μg.mg^−1^ DW) are similar both on green and on yellow leaves (Supplement 2, middle column), whereas in leaf tissues mined by tissue-feeders, starch quantities (*green leaves*: 77.31 ± 5.10 μg.mg^−1^ DW; *yellow leaves*: 48.76 ± 5.01 μg.mg^−1^ DW) differ on green and on yellow leaves (Supplement 2A, right column) and are close to their respective controls (Supplement 2C).

#### Amino acids

Senescence leads to a metabolic reconfiguration of the leaves with a drastic reduction of the protein-bound amino acid content (from 1.98 ± 0.20 to 0.65 ± 0.06 μg.mg^−1^ DW) and an increase of free amino acid content (from 0.17 ± 0.01 to 0.39 ± 0.03 μg.mg^−1^ DW) (Figures 3, 4 and 5; Supplements 3 and 4, left columns). In mined tissues, both on green and on yellow leaves, the protein-bound amino acid content remains stable and similar to unmined green controls (Figures 3A and 3B; Supplement 3A, middle and right columns; Supplements 3B and 3C, left columns; Supplement 3D). This pattern can be observed for both fluid- and tissues-feeders and for almost each individual protein-bound amino acid (Figure 4). Contrary to protein-bound amino acids, free amino acid content is closely associated to larvae developmental stages (Figures 3C and 3D; Supplement 4A, middle and right columns; Supplements 4B, 4C and 4D) with pattern observed in leaf tissues mined by tissue-feeding larvae being different from fluid-feeders. Indeed, leaf tissues mined by fluid-feeding larvae show a strong increase of free amino acids on yellow leaves whereas the level remains low on green leaves (Figure 3C; Supplement 4A, middle column; Supplement 4B). However, tissue-feeders induce a strong increase of free amino acid content in mined tissues both on green and on yellow leaves (Figure 3D; Supplement 4C). As a consequence, amino acid content of mined tissues differs between green and yellow leaves for fluid-feeding larvae (Figure 3C; Supplement 4A, middle column), but is similar on green and on yellow leaves for tissue-feeding larvae (Figure 3D; Supplement 4A, right columns). Changes observed in free amino acid content in mined tissues are mostly due to alteration of non-essential amino acid concentrations (Figure 3C). In tissues mined by fluid-feeders, the strong increase of the most abundant non-essential free amino acid, glutamine (50 %) (and asparagine (10 %) to a lesser extent) is responsible for these changes (Figure 5). Moreover, apple-tree leaves appear to provide very low amounts of methionine (~0.5 % of total amino acids), histidine (~1 %), and tryptophan (~1 %).

### Essential amino acid composition of the leaf-mining larva and limiting amino acids

The whole-body amino acid composition (Table 1) of *P. blancardella* larvae was used to estimate their essential amino acid requirements. It appears that larvae have a strong demand in histidine (EAA; ~50 % of their amino acid pool; Table 1A) for both larval feeding modes, in cysteine for fluid-feeders (NEAA; ~20 %; Table 1B), and in lysine for tissue-feeders (EAA; ~12 %; Table 1A).

**Table 1.** (**A**) Essential and (**B**) non-essential amino acid (protein-bound + free) content of whole body of fluid- and tissue-feeding leaf-mining insects both on green and on yellow leaves. Data are presented as average ± S.E.M. and expressed in μg per mg of caterpillar dry weight. Values in rows with different letters (a, b) are significantly different (P < 0.05).

**A. Essential amino acids**
|  | Green leaves |  | Yellow leaves |  |
| --- | --- | --- | --- | --- |
|  | Fluid-feeders | Tissue-feeders | Fluid-feeders | Tissue-feeders |
| Val | 12.48 ± 0.67 a | 10.39 ± 1.69 a | 10.75 ± 1.78 a | 8.74 ± 1.55 a |
| Leu | 25.72 ± 1.42 a | 13.18 ± 2.23 b | 20.64 ± 5.92 a,b | 11.65 ± 2.58 a,b |
| Ile | 8.98 ± 0.83 a | 3.73 ± 0.71 b | 9.84 ± 2.33 a,b | 3.48 ± 0.75 a,b |
| Thr (+ Ser) | 2.52 ± 0.93 a | 1.46 ± 0.29 a | 2.57 ± 0.90 a | 1.33 ± 0.24 a |
| Arg (+ Asp) | 18.15 ± 2.75 a | 14.29 ± 1.94 a | 19.62 ± 1.23 a | 12.03 ± 1.59 a |
| Met | 2.01 ± 0.29 a | 0.54 ± 0.08 b | 5.65 ± 1.60 c | 0.48 ± 0.12 b |
| Phe (+ Glu) | 12.75 ± 0.89 a | 13.47 ± 1.84 a | 7.65 ± 0.89 a | 12.63 ± 2.01 a |
| Lys | 14.95 ± 1.78 a | 54.48 ± 9.49 b | 10.96 ± 6.31 a | 42.27 ± 5.86 b |
| His | 234.06 ± 45.73 a | 140.67 ± 45.64 a | 464.42 ± 204.74 a | 243.82 ± 112.25 a |
| Trp | 1.63 ± 0.55 a | 11.88 ± 9.79 a | 3.70 ± 1.53 a | 1.01 ± 0.29 a |

**B. Non-essential amino acids**
|  | Green leaves |  | Yellow leaves |  |
| --- | --- | --- | --- | --- |
|  | Fluid-feeders | Tissue-feeders | Fluid-feeders | Tissue-feeders |
| Ala | 5.35 ± 0.88 a | 5.07 ± 0.94 a | 7.07 ± 0.81 a | 3.96 ± 0.67 a |
| Gly | 11.80 ± 0.63 a | 8.68 ± 1.52 a | 9.02 ± 1.82 a | 8.00 ± 1.55 a |
| Pro | 11.81 ± 0.21 a | 8.91 ± 1.32 a | 7.64 ± 0.58 a | 8.01 ± 1.20 a |
| Asn | 7.15 ± 1.79 a | 1.28 ± 0.45 b | 10.00 ± 2.43 a | 1.72 ± 0.75 b |
| Cys | 88.88 ± 8.83 a | 24.43 ± 9.55 b | 180.52 ± 59.49 a | 24.71 ± 9.88 b |
| Gln | 2.72 ± 0.33 a | 1.38 ± 0.35 a | 4.55 ± 1.53 a | 1.29 ± 0.42 a |
| Tyr | 1.86 ± 0.62 a | 15.84 ± 4.95 a | 1.60 ± 0.64 a | 6.11 ± 2.52 a |

The relative limitation of EAAs was quantified with chemical scores, with the lowest scores indicating that His was the first most limiting EAA for both fluid- and tissue feeding instars and both on green and on yellow leaves (chemical scores between 0.01 and 0.05) (Table 2). Therefore, His would be the first EAA to be depleted during protein synthesis in *P. blancardella*.

**Table 2.**
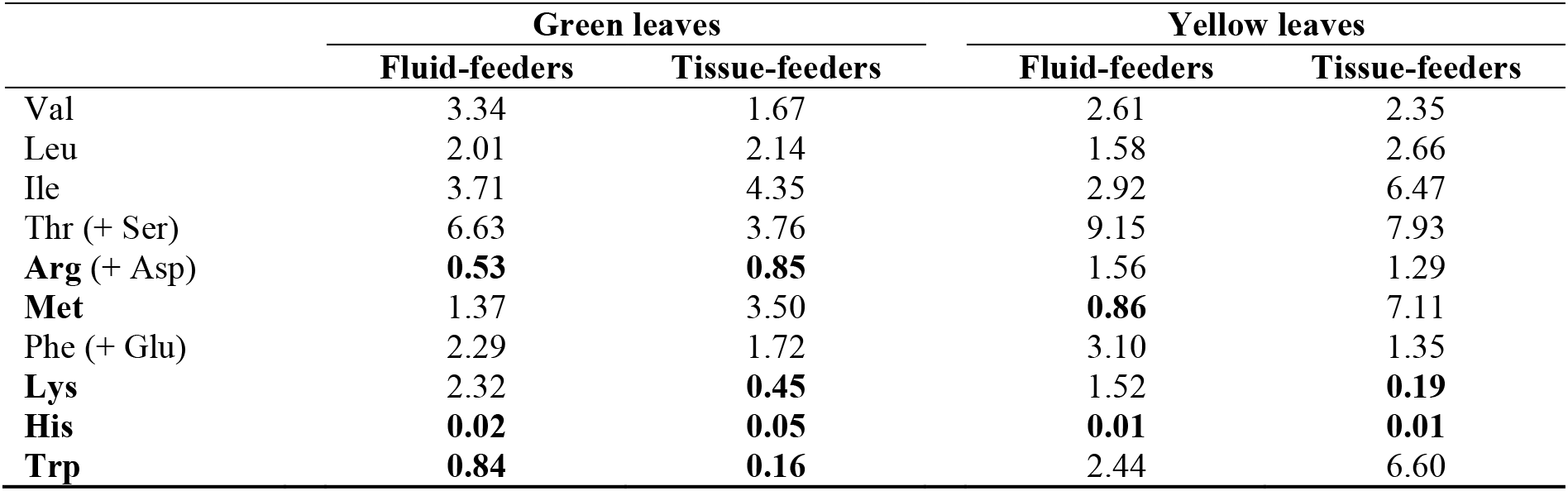
Chemical scores of essential amino acids (EAAs) in mined tissues both on green and on yellow leaves based on the EAA compositions of the whole body of *Phyllonorycter blancardella* leaf-miner. When EAAs were co-eluted with NEAAs, the name of the NEAA is in brackets. Chemical scores were calculated as (μg of EAA in mined leaf tissues / μg of total AA in mined leaf tissues) / (μg of EAA in insect/μg of total AA in insect). Limiting essential amino acids are in bold for both fluid- and tissue-feeding instars. A value lower than 1 indicates an excess of EAA available in the foliar amino acid pool. *Val* valine, *Leu* leucine, *Ile* isoleucine, *Thr* (+ *Ser*) threonine (+ serine), *Arg* (+ *Asp*) arginine (+ aspartate), *Met* methionine, *Phe* (+ *Glu*) phenylalanine (+ glutamate), *Lys* lysine, *His* histidine, *Trp* tryptophan.

|  | Green leaves |  | Yellow leaves |  |
| --- | --- | --- | --- | --- |
|  | Fluid-feeders | Tissue-feeders | Fluid-feeders | Tissue-feeders |
| Val | 3.34 | 1.67 | 2.61 | 2.35 |
| Leu | 2.01 | 2.14 | 1.58 | 2.66 |
| Ile | 3.71 | 4.35 | 2.92 | 6.47 |
| Thr (+ Ser) | 6.63 | 3.76 | 9.15 | 7.93 |
| **Arg** (+ Asp) | **0.53** | **0.85** | 1.56 | 1.29 |
| **Met** | 1.37 | 3.50 | **0.86** | 7.11 |
| Phe (+ Glu) | 2.29 | 1.72 | 3.10 | 1.35 |
| **Lys** | 2.32 | **0.45** | 1.52 | **0.19** |
| **His** | **0.02** | **0.05** | **0.01** | **0.01** |
| **Trp** | **0.84** | **0.16** | 2.44 | 6.60 |

Moreover, senescence induces changes in amino acid limitation. Indeed, arginine (+ aspartate) and tryptophan appear to be limiting only for larvae developing on green leaves, primarily due to their higher availability on yellow leaves (Figures 5.8 and 5.14; Table 2). It is also important to note changes in amino acid requirements according to the development stage of larvae. Tissue-feeding larvae are indeed limited by lysine whereas fluid-feeders are not (Table 2). This limitation is mostly due to an increased demand for this amino acid in later stages (Table 1A). Moreover, methionine seems to be limiting only for fluid-feeders developing on yellow leaves (Table 2). Finally, on yellow leaves, the second most limiting EAA is less limiting for fluid-feeders (0.86 for methionine) than for tissue-feeders (0.19 for lysine) (Table 2). More generally, *P. blancardella* larvae living on yellow leaves appear to be less limited (2 limiting EAAs) than larvae on green leaves (3-4 limiting EAAs) (Table 2).

## Discussion

### Strong alteration of plant primary metabolism in mined tissues – Are they becoming functionally independent areas?

According to our predictions, primary metabolism was altered in the tissue mined by the feeding activity of the insect, with a modification of the total amount of sugars, the specific sugar composition and the protein-bound and free amino acid contents. We hypothesized that mines would act as active nutrient sinks and preferentially accumulate nutrients (Schwachtje and Baldwin, 2008; Bolton, 2009; Kerchev et al., 2012). However, the nutritional content of mined tissues was indeed modified, but in the opposite direction. Sugars and protein-bound amino acids are less (fluid-feeders, for sugars only) or equally (tissue-feeders) concentrated in mined tissues both on green and on yellow leaves than in unmined green controls (Figures 1, 3, 4 and 5). Free amino acids, by contrast, accumulate into mined tissues as predicted.

Our results show that protein composition in mined tissues does not seem to be altered neither as part of the senescing process, as a plant defensive mechanism, nor as a leaf manipulation by the insect for its own benefit. Indeed, the protein-bound amino acid composition experienced by caterpillars feeding on senescing leaves (i.e. mined tissues displaying a green-island) is close to the composition of mined tissues on green leaves, and mined tissues both on green and on yellow leaves have a similar composition to unmined green control tissues (Figures 3 and 4). Free amino acid levels, by contrast, are altered in mined tissues with a pattern closely associated with larval development stages. Free amino acid content is higher in tissues mined by tissue-feeding instars both on green and on yellow leaves, whereas this increase is greater and only visible on yellow leaves for fluid-feeding instars. This change of free amino acid content observed for younger larvae on senescing leaves is essentially due to a strong increase of glutamine. For later instars, it is mainly due to an increase of glycine, arginine + aspartate, asparagine and glutamine (Figures 3 and 5).

Our results thus strengthen the hypothesis that mines are functionally independent areas, operating a metabolic machinery of their own and providing a “dietary buffer” to the insect, especially in an otherwise senescent autumnal environment (Engelbrecht et al., 1969; Giron et al., 2007, Body et al., 2013). The higher amounts of sugar in mines inhabited by tissue-feeding larvae on yellow leaves (compared to unmined yellow controls; Supplement 2C, right column) are not the outcome of an accumulation process, but of a localized continuous renewal. The absence of remobilization of sugars and amino acids by the plant from the insect’s feeding area in the Fall (similar nutrient content in unmined zones UM^1^, UM^2^ and UM^3^) and the similar composition of mined tissues both on green and on yellow leaves strongly reinforce the concept of nutritional homeostasis within mined tissues. Similar results have been found in the gall-inducing insects *Neuroterus quercus-baccarum* and *Andricus lignicola* for which induced gall tissues contain less nutrient than surrounding tissues (Hartley and Lawton, 1992; Diamond et al., 2008).

### Insect manipulation rather than plant defense

Nutritional requirements of any leaf-mining insect were completely unknown before this study, and the lack of artificial diets, preventing manipulative experiments, is a real bottleneck. Changes in plant primary metabolism after biotic infestations have often been interpreted as a necessary requirement to satisfy the increased demand for energy and carbon skeletons to sustain the direct defense machinery and corresponding physiological adaptations (Kerchev et al., 2012). It is also suggested that primary metabolites could function as signaling molecules in plant defense pathways or could act as direct plant defensive compounds (Augner, 1995; Berenbaum, 1995; Schwachtje and Baldwin, 2008). Primary metabolism reconfiguration could also potentially allow the plant to tolerate herbivory while minimizing impacts on fitness traits by supporting necessary physiological adjustments plants must make (Fornoni, 2011). Changes observed in amino acid abundance may not directly impact larval nutrition, but may also be associated with the changing physiological process in the host-plant as it senesces and/or adapts to the leaf-mining larvae (plant tolerance or defense). Indeed, asparagine and glutamine, major components of the free amino acid pool (Figure 5), are known as important nitrogen carriers in plants and involved in remobilization of leaf nitrogen during leaf senescence or infection by pathogens (Lea and Miflin, 1980; Sieciechowicz et al., 1988; Lam et al., 1996; Lea et al., 2007). When plant leaves are attacked by fungi or bacteria, asparagine levels rapidly increase in infected leaves (Sieciechowicz et al., 1988; Pérez-García et al., 1998; Scarpari et al., 2005). An increase in the level of asparagine is usually interpreted as a plant reaction against the pathogen due to recovery of leaf protein from attacked leaves (Pérez-García et al., 1998; Scarpari et al., 2005). Insect growth may also strongly depend on the effects of secondary plant substances and/or carbon:nitrogen ratios in the diet (Berenbaum, 1995; Schoonhoven et al., 2005). Thus, even though the physiological function of altered amino acids in mined tissues remains unclear, our results suggest that such alterations of leaf amino acid profiles contribute to enhance the nutritional quality of plant tissues ingested by larvae and may specifically contribute to increase larval fitness under senescing conditions (Table 2). Additionally, in the insect’s feeding area, the concomitant inhibition of classical plant defenses (plant secondary compounds such as phenolics – Giron et al., 2016) also strongly suggest the ability of leaf-mining insects to manipulate their host-plant to improve their nutritional environment, leading to a high survival rate in the absence of natural enemies (90 %; Pottinger and LeRoux, 1971; Giron, personal communication).

### Mechanisms of plant manipulation

The morphological impact of larvae on leaf tissues could be, at least partially, responsible for the differential modulation of the leaf physiological response according the larval feeding-mode. Such differential control of plant sugars (Figure 2, pie chart #4) could also potentially be explained by changes in insect saliva composition (including level and composition of cytokinins; Zhang et al., 2017) and/or in endosymbiotic bacteria levels over the course of the insect development. Indeed, *P. blancardella* and other Gracillariidae moths prevent mined tissues from senescing (inducing a “green-island” phenotype) through endosymbiotic bacteria mediated release of cytokinins which are known to positively impact plant sugar metabolism (Giron et al., 2007; Kaiser et al., 2010; Body et al., 2013; Gutzwiller et al., 2015; Zhang et al., 2017). *P. blancardella* seems thus to take advantage of its intimate association with the plant by controlling nutrient levels in mined tissues for its own benefit (Body et al., 2013; this study). This is consistent with previous results showing the capacity of this insect to alter the plant hormonal balance (Engelbrecht et al., 1969; Giron et al., 2007; Kaiser et al., 2010; Body et al., 2013; Zhang et al., 2016), directly impacting plant primary metabolism (Ehneß and Roitsch, 1997; Lara et al., 2004; Roitsch and González, 2004; Walters and McRoberts, 2006).

### Leaf-mining larvae make the best out of scenescing plant tissues

For *P. blancardella*, the fitness consequences of the sugar content regulation, allowing survival under adverse conditions to complete an additional generation in the Fall, are significant (Kaiser et al., 2010; Body et al., 2013). However, fluid-feeding instars appear to partially suffer from leaf senescence due to their limited abilities to control the sugar content of mined tissues on yellow leaves (Figure 2, pie chart #4; Supplement 2B, right column, sugar composition). In fact, by closely estimating qualitative and quantitative sugar composition of leaf tissues for the two distinct larval stages, we show that the nutritional landscape experienced by fluid-feeding caterpillars feeding on senescing leaves is not dissimilar to unmined yellow leaves, indicative of a lack of control by the fluid-feeding larvae. By contrast, tissue-feeding larvae have acquired extended capacities to regulate the sugar content (for both sugar quantities and composition) in order to ‘delay’ the leaf senescing process (Figure 2, scenarii B and D; Supplement 2C) and eventually sometimes ‘reversing’ the senescing process already engaged (Figure 2, scenarii C and E; Supplements 2B vs. 2C, right columns, sugar composition). This allows insects to generate a nutritional environment similar to green leaves for sugars (except for fluid-feeders on yellow leaves; Figure 2, pie chart #4). Analyzing specific sugars, one notes that sorbitol represents a qualitatively and quantitatively important part of the sugar content in mined tissues (Figures 1 and 2). Its up-regulation on yellow leaves for tissue-feeders may allow larvae to overcome freezing in late Fall. Sugar alcohols are indeed often involved in thermal tolerance, particularly in cryoprotection (Sømme, 1965; Wyatt, 1967; Miller and Smith, 1975; Wolfe et al., 1998; Salvucci et al., 2000). In summary, the manipulation of sugar content at the feeding site most likely allows for better insect performances, as observed for the forest tent caterpillar *Malacosoma disstria* for which a sugar regulation increases caterpillar survival rate (Noseworthy and Despland, 2006).

### Beyond leaf-miners

While fitness consequences of an increase availability of amino acids for *P. blancardella* are still unknow, a recent study on another leaf-miner, *Tuta absoluta,* showed that nutrition on nitrogen-deficient plant tissues impaired the leaf-miner development by notably decreasing pupal weight and lengthening the development period from egg to adult (Mattson, 1980; Larbat et al., 2016). Transposed to our system, a nitrogen-enriched food source on yellow leaves could thus favor a faster development to reach the tissue-feeding stage before climatic conditions become detrimental for insect survival (Pottinger and LeRoux, 1971; Edwards and Wratten, 1980). Some leaf-miners are able to change from one mine (or one leaf) to another in case of inadequate food supply (Needham et al., 1928) – such as the Diptera larva *Scaptomyza flava* (Whiteman et al., 2011), the Coleoptera larva *Neomycta rubida* (Martin, 2010) and the micro-Lepidoptera *Coleophora klimeschiella* (Khan and Baloch, 1976) – but such strategies are very rare and only restricted to certain groups. Unlike these temporary miners, *P. blancardella* larvae are constrained to their mine.

The lack of comparative work on the EAA requirements of herbivores, and more specifically of leaf-feeding insects, and the amino acid composition of the plant consumed (see for an exception: Barbehenn et al., 2013) has hampered the understanding of many plant-herbivore interactions, with implications ranging from insect behavioral ecology to adaptation, speciation, and population dynamics. In our system, the use of chemical scores allowed us to show that leaf-mining larvae experience a lower EAA limitation when feeding on yellow leaves than on green leaves. Improved nitrogen availability is thus experienced by these insects during the senescence process. Our results are consistent with other studies that have identified histidine, methionine and arginine as the most limiting amino acids for herbivore species. Histidine and methionine were indeed the first or second most limiting EAAs for the caterpillar *Lymantria dispar* throughout the growing season (Barbehenn et al., 2013). Several vertebrate herbivores follow the same pattern. Springboks (*Antidorcas marsupialis*), blesboks (*Damaliscus dorcas phillipsi*) and impalas (*Aepyceros melampus*) have also been shown to be limited by these same three amino acids: methionine, histidine, and arginine (Van Zyl and Ferreira, 2003). These similarities between caterpillars and large herbivores suggest that these amino acid limitations are a direct consequence of using plants as food source in general.

## Synthesis

### Nutritional intimate interactions between a leaf-miner and its senescing leaf

The following putative scheme synthesizes the observed sequence of mechanisms at play in the interaction between a leaf-miner and its leaf during Fall. First, symbiont-mediated increase of cytokinin profiles would induce an accumulation of sugar at the leaf-miner feeding site (Giron et al., 2007; Kaiser et al., 2010; Body et al., 2013; Zhang et al., 2016, 2017; this study). Then, to maintain nutritional homeostasis at the feeding site, carbohydrates in excess would be instead degraded by the plant to fuel the amino acid biosynthetic pathways. Indeed, they are costly to process for the larva and could bride sugar and amino acid metabolisms. This is a key assumption, requiring further investigation on biochemical pathways in this biological system. However, such degradation of excess carbohydrates and their conversion into intermediates for amino acid synthesis has been observed for the Hessian fly *M. destructor* (Liu et al., 2007). The observed partial manipulation of plant tissue nutritional value mitigates the detrimental effects of senescent leaves as food source for *P. blancardella* in the race against time for completing an extra-generation.

## Acknowledgements

We would like to thank Alexandre Lebrun, Isabelle Mazerie, and Jean-Philippe Christidès for their help with sugars and amino acids methods and analyses. We would also like to thank Wilfried Kaiser, Elisabeh Huguet, Géraldine Dubreuil, Jean-Paul Monge, Arnaud Lanoue, Gaëlle Glevarec, Benoît St-Pierre, Vincent Courdavault, and the “Endofeed team” for helpful discussions, as well as Laurent Ardouin for free access to his orchard. This study was supported by the ANR project ECOREN ANR-JC05-46491 and the Région Centre project 201000047141 to D. Giron.

## References

Anderson, T.R., Boersma, M., Raubenheimer, D. (2004). Stoichiometry: Linking elements to biochemicals. Ecol. 85, 1193–1202. doi: 10.1890/02-0252

Augner, M. (1995). Low nutritive quality as a plant defence: Effects of herbivore-mediated interactions. Evol. Ecol, 9, 605–616. doi: 10.1007/BF01237658

Barbehenn, R.V., Niewiadomski, J., Kochmanski, J. (2013). Importance of protein quality versus quantity in alternative host plants for a leaf-feeding insect. Oecol. 173, 1–12. doi: 10.1007/s00442-012-2574-7

Berenbaum, MR. (1995). Turnabout is fair play: Secondary roles for primary compounds. J Chem. Ecol. 21, 925–940. doi: 10.1007/BF02033799

Bernays, E.A., Chapman, R.F. (1994). Host-plant selection by phytophagous insects. Chapman and Hall, New York.

Body, M., Kaiser, W., Dubreuil, G., Casas, J., Giron, D. (2013). Leaf-miners co-opt microorganisms to enhance their nutritional environment. J. Chem. Ecol. 39, 969–977. doi: 10.1007/s10886-013-0307-y

Body, M., Burlat, V., Giron, D. (2015). Hypermetamorphosis in a leaf-miner allows insects to cope with a confined nutritional space. Arthropod-Plant Interact. 9, 75–84. doi: 10.1007/s11829-014-9349-5

Body, M., Casas, J., Christidès, J.-P., Giron, D. (2018). Underestimation of carbohydrates by classical anthrone-based colorimetric techniques compromises insect metabolic and energetic studies. BioRXiv. doi: 10.1101/322123

Bolton, M.D. (2009). Primary metabolism and plant defense: Fuel for the fire. Mol. Plant-Microbe Interact. 22, 487–497. doi: 10.1094/MPMI-22-5-0487

Chiou, S.H., Wang, K.T. (1988). Simplified protein hydrolysis with methanesulphonic acid at elevated temperature for the complete amino acid analysis of proteins. J. Chromatogr. A. 448, 404–410.

Chown, S.L., Nicolson, S.W. (2004). Insect physiological ecology, mechanisms and patterns. Oxford University Press.

Connor, E.F., Taverner, M.P. (1997). The evolution and adaptive significance of the leaf-mining habit. Oikos 79, 6625–6645. doi: 10.2307/3546085

Dadd, R.H. (1985). Nutrition: Organisms. Comparative Insect Physiology, Biochemistry, and Pharmacology 4, 313–390.

Dardeau, F., Body, M., Berthier, A., Miard, F., Christidès, J.-P., Feinard-Duranceau, M., Brignolas, F., Giron, D., Lieutier, F., Sallé, A. (2015). Effects of fertilisation on amino acid mobilisation by a plant-manipulating insect. Ecol. Entomol. 40, 814–822. doi: 10.1111/een.12274

Diamond, S.E., Blair, C.P., Abrahamson, W.G. (2008). Testing the nutrition hypothesis for the adaptative nature of insect galls: Does a non-adapted herbivore perform better in galls? Ecol. Entomol. 33, 385–393. doi: 10.1111/j.1365-2311.2007.00979.x

Douglas, A.E. (2006). Phloem sap feeding by animals: Problems and solutions. J. Exp. Bot. 57, 747–754. doi: 10.1093/jxb/erj067

Douglas, A.E. (2009). The microbial dimension in insect nutritional ecology. Funct. Ecol. 23, 38–47. doi: 10.1111/j.1365-2435.2008.01442.x

Douglas, A.E. (2013). Microbial brokers of insect-plant interactions revisited. J. Chem. Ecol. 39, 952–961. doi: 10.1007/s10886-013-0308-x

Douglas, A.E., Minto, L.B., Wilkinson, T.L. (2001). Quantifying nutrient production by the microbial symbiosis in an aphid. J. Exp. Biol. 204, 349–358.

Edwards, P.J., Wratten, S.D. (1980). “Chapter 2: The problems of plant as food for animals”, in Ecology of insect-plant interactions, ed. E. Arnold. The Camelot Press Ltd, Southampton. Studies in Biology no. 121.

Ehneß, R., Roitsch, T. (1997). Co-ordinated induction of mRNAs for extracellular invertase and a glucose transporter in *Chenopodium rubrum* by cytokinins. Plant J. 11, 539–548. doi: 10.1046/j.1365-313X.1997.11030539.x

Eleftherianos, I., Vamvatsikos, P., Ward, D., Gravanis, F. (2006). Changes in the levels of plant total phenols and free amino acids induced by two cereal aphids and effects on aphid fecundity. J. Appl. Entomol. 130, 15–19. doi: 10.1111/j.1439-0418.2005.01017.x

Engelbrecht, L., Orban, U., Heese, W. (1969). Leaf-miner caterpillars and cytokinins in the green islands of autumn leaves. Nature 223, 319–321. doi: 10.1038/223319a0

Fornoni, J. (2011). Ecological and evolutionary implications of plant tolerance to herbivory. Funct. Ecol. 25, 399–407. doi: 10.1111/j.1365-2435.2010.01805.x

Fountoulakis, M., Lahm, H.W. (1998). Hydrolysis and amino acid composition analysis of proteins. J. Chromatogr. A 826, 109–134. doi: 10.1016/S0021-9673(98)00721-3

Giron, D., Glevarec, G. (2014). Cytokinin-induced phenotypes in plant-insect interactions: Learning from the bacterial world. J. Chem. Ecol. 40, 826–835. doi: 10.1007/s10886-014-0466-5

Giron, D., Rivero, A., Mandon, N., Darrouzet, E., Casas, J. (2002). The physiology of host feeding in parasitic wasps: Implications for survival. Funct. Ecol. 16, 750–757. doi: 10.1046/j.1365-2435.2002.00679.x

Giron, D., Kaiser, W., Imbault, N., Casas, J. (2007). Cytokinin-mediated leaf manipulation by a leaf-miner caterpillar. Biol. Lett. 3, 340–343. doi: 10.1098/rsbl.2007.0051

Giron, D., Huguet, E., Stone, G.N., Body, M. (2016). Insect-induced effects on plants and possible effectors used by galling and leaf-mining insects to manipulate their host-plant. J. Insect Physiol. 84, 70–89. doi: 10.1016/j.jinsphys.2015.12.009

Gündüz, E.A., Douglas, A.E. (2009). Symbiotic bacteria enable insect to use a nutritionally inadequate diet. Proc. Roy. Soc. B – Biol. Sci. 279, 987–991. doi: 10.1098/rspb.2008.1476

Gutzwiller, F., Dedeine, F., Kaiser, W., Giron, D., Lopez-Vaamonde, C. (2015). Correlation between the green-island phenotype and *Wolbachia* infections during the evolutionary diversification of Gracillariidae leaf-mining moths. Ecol. Evol. 5, 4049–4062. doi: 10.1002/ece3.1580

Hansen, J., Møller, I. (1975). Percolation of starch and soluble carbohydrates from plant tissue for quantitative determination with anthrone. Anal. Biochem. 68, 87–94. doi: 10.1016/0003-2697(75)90682-X

Hartley, S.E. (1998). The chemical composition of plant galls: Are levels of nutrients and secondary compounds controlled by the gall-former? Oecol. 113, 492–501. doi: 10.1007/s004420050401

Hartley, S.E., Lawton, J.H. (1992). Host-plant manipulation by gall-insects: A test of the nutrition hypothesis. J. Anim. Ecol. 61, 113–119. doi: 10.2307/5514

Hering, M.E. (1951). “Chapter 5: Changing from one mine to another”, in Biology of the leaf-miner, ed. W. Junk. Gravenhage, Berlin, 33–38.

Horie, Y., Watanabe, K. (1983). Effect of various kinds of dietary protein and supplementation with limiting amino acids on growth, haemolymph components and uric acid excretion in the silkworm, *Bombyx mori*. J. Insect Physiol. 29, 187–199. doi: 10.1016/0022-1910(83)90143-9

Kaiser, W., Huguet, E., Casas, J., Commin, C., Giron, D. (2010). Plant green-island phenotype induced by leaf-miners is mediated by bacterial symbionts. Proc. Roy. Soc. B – Biol. Sci. 277, 2311–2319. doi: 10.1098/rspb.2010.0214

Karasov, W.H., Martinez Del Rio, C. (2007). Physiological ecology: How animals process energy, nutrients, and toxins. Princeton University Press, Princeton.

Karley, A.J., Douglas, A.E., Parker, W.E. (2002). Amino acid composition and nutritional quality of potato leaf phloem sap for aphids. J. Exp. Biol. 205, 3009–3018.

Karowe, D.N., Martin, M.M. (1989). The effects of quantity and quality of diet nitrogen on the growth, efficiency of food utilization, nitrogen budget, and metabolic rate of fifth-instar *Spodoptera eridania* larvae (Lepidoptera: Noctuidae). J. Insect Physiol. 35, 699–708. doi: 10.1016/0022-1910(89)90089-9

Kerchev, P.I., Fenton, B., Foyer, C.H., Hancock, R.D. (2012). Plant responses to insect herbivory: Interactions between photosynthesis, reactive oxygen species and hormonal signaling pathways. Plant Cell Environ. 35, 441–453. doi: 10.1111/j.1365-3040.2011.02399.x

Kessler, A., Baldwin, I.T. (2002). Plant responses to insect herbivory: The emerging molecular analysis. Annu. Rev. Plant Biol. 53, 299–328. doi: 10.1146/annurev.arplant.53.100301.135207

Khan, A.G., Baloch, G.M. (1976). *Coleophora klimeschiella* [Lep; Coleophoridae] a promising biocontrol agent for Russian thistles, *Salsola spp*. Entomophaga 21, 425–428. doi: 10.1007/BF02371641

Kimmerer, T.W., Potter, D.A. (1987). Nutritional quality of specific leaf tissues and selective feeding by a specialist leaf-miner. Oecol. 71, 548–551. doi: 10.1007/BF00379295

Klowden, M.J. (2007). Physiological systems in insects. Second edition. Elsevier.

Koyama, Y., Yao, I., Akimoto, SI. (2004). Aphid galls accumulate high concentrations of amino acids: A support for the nutrition hypothesis for gall formation. Entomol. Exp. Appl. 113, 35–44. doi: 10.1111/j.0013-8703.2004.00207.x

Lam, H.M., Coschigano, K.T., Oliveira, I.C., Melo-Oliveira, R., Coruzzi, G.M. (1996). The molecular genetics of nitrogen assimilation into amino acids in higher plants. Annu. Rev. Plant Biol. 47, 569–593. doi: 10.1146/annurev.arplant.47.1.569

Lara, M.E.B., Garcia, M.C.G., Fatima, T., Ehneß, R., Lee, T.K., Proels, R., Tanner, W., Roitsch, T. (2004). Extracellular invertase is an essential component of cytokinin-mediated delay of senescence. Plant Cell 16, 1276–1287. doi: 10.1105/tpc.018929

Larbat, R., Adamowicz, S., Robin, C., Han, P., Desneux, N., Le Bot, J. (2016). Interrelated responses of tomato plants and the leaf miner *Tuta absoluta* to nitrogen supply. Plant Biol. 18, 495–504. doi: 10.1111/plb.12425

Larson, K.C., Whitham, T.G. (1991). Manipulation of food resources by a gall-forming aphid: The physiology of sink-source interactions. Oecol. 88, 15–21. doi: 10.1007/BF00328398

Lea, P.J., Miflin, B. (1980). “Transport and metabolism of asparagine and other nitrogen compounds within the plant”, in The biochemistry of plants: Amino acids and derivatives, ed. B.J. Milflin. Academic, New York, 569–607.

Lea, P.J., Sodek, L., Parry, M.A.J., Shewry, P.R., Halford, N.G. (2007). Asparagine in plants. Ann. Appl. Biol. 150, 1–26. doi: 10.1111/j.1744-7348.2006.00104.x

Liu, X., Bai, J., Huang, L., Zhu, L., Liu, X., Weng, N., Reese, J.C., Harris, M.O., Stuart, J.J., Chen, M.S. (2007). Gene expression of different wheat genotypes during attack by virulent and avirulent Hessian fly (*Mayetiola destructor*) larvae. J. Chem. Ecol. 33, 2171–2194. doi: 10.1007/s10886-007-9382-2

Machado, R.A., Ferrieri, A.P., Robert, C.A.M., Glauser, G., Kallenbach, M., Baldwin, I.T., Erb, M. (2013). Leaf-herbivore attack reduces carbon reserves and regrowth from the roots via jasmonate and auxin signaling. New Phytol. 200, 1234–1246. doi: 10.1111/nph.12438

Machado, R.A., Arce, C., Ferrieri, A.P., Baldwin, I.T., Erb, M. (2015). Jasmonate-dependent depletion of soluble sugars compromises plant resistance to *Manduca sexta*. New Phytol. 207, 91–105. doi: 10.1111/nph.13337

Marshall, J.D. (1986). Drought and shade interact to cause fine-root mortality in Douglas-fir seedlings. Plant Soil 91, 51–60. doi: 10.1007/BF02181818

Martin, N.A. (2010). Pohutakawa leaf miner – *Neomycta rubida*. New Zealand Arthropod Collection Factsheet Serie.

Mattson, W.J. Jr. (1980). Herbivory in relation to plant nitrogen content. Annu. Rev. Ecol. Syst. 11, 119–161.

Miller, L.K., Smith, J.S. (1975). Production of threitol and sorbitol by an adult insect: Association with freezing tolerance. Nature 258, 519–520. doi: 10.1038/258519a0

Nakamura, M., Miyamoto, Y., Ohgushi, Y. (2003). Gall initiation enhances the availability of food resources for herbivorous insects. Funct. Ecol. 17, 851–857. doi: 10.1111/j.1365-2435.2003.00786.x

Nation, J.L. (2002). “Chapter 3: Nutrition”, in Insect Physiology and Biochemistry, ed. J.L. Nation. CRC Press, 65–88.

Needham, J.G., Frost, S.W., Tothill, B.H. (1928). “Chapter 2: Extend of the leaf-mining habit”, in Leaf-mining insects. Ecology Classics. Williams and Wilkins Company, Baltimore, MD, 36–40.

Noseworthy, M.K., Despland, E. (2006). How do primary nutrients affect the performance and preference of forest tent caterpillars on trembling aspen? Can. Entomol. 138, 367–375. doi: 10.4039/n05-076

Oren, R., Schulze, E.D., Werk, K.S., Meyer, J., Schneider, B.U., Heilmeier, H. (1988). Performance of two *Picea abies* (L.) Karst. stands at different stages of decline. Oecol. 75, 25–37. doi: 10.1007/BF00378810

Pérez-García, A., Pereira, S., Pissarra, J., García Gutiérrez, A., Cazorla, F.M., Salema, R., De Vicente, A., Cánovas, F.M. (1998). Cytosolic localization in tomato mesophyll cells of a novel glutamine synthetase induced in response to bacterial infection or phosphinothricin treatment. Planta 206, 426–434. doi: 10.1007/s004250050418

Pottinger, R.P., LeRoux, E.J. (1971). The biology and dynamics of *Lithocolletis blancardella* (Lepidoptera: Gracillariidae) on apple in Quebec. Memoirs of the Entomological Society of Canada no. 77. doi: 10.4039/entm10377fv

Rock, G.C. (1972). “Optimal proportions of dietary amino acids”, in Insect and mite nutrition, ed. J.G. Rodriguez. North-Holland, Amsterdam, 183–197.

Roitsch, T., González, M.C. (2004). Function and regulation of plant invertases: Sweet sensations. Trends Plant Sci. 9, 606–613. doi: 10.1016/j.tplants.2004.10.009

Rovio, S., Yli-Kauhaluoma, J., Sirén, H. (2007). Determination of neutral carbohydrates by CZE with direct UV detection. Electrophoresis 28, 3129–3135. doi: 10.1002/elps.200600783

Saltzmann, K.D., Giovanini, M.P., Zheng, C., Williams, C.E. (2008). Virulent Hessian fly larvae manipulate the free amino acid content of host wheat plants. J. Chem. Ecol. 34, 1401–1410. doi: 10.1007/s10886-008-9544-x

Salvucci, M.E., Stecher, D.S., Henneberry, T.J. (2000). Heat shock proteins in whiteflies, an insect that accumulates sorbitol in response to heat stress. J. Therm. Biol. 25, 363–371. doi: 10.1016/S0306-4565(99)00108-4

Scarpari, L.M., Meinhardt, L.W., Mazzafera, P., Pomella, A.W.V., Schiavinato, M.A., Cascardo, J.C.M., Pereira, G.A.G. (2005). Biochemical changes during the development of witches’ broom: The most important disease of cocoa in Brazil caused by *Crinipellis perniciosa*. J. Exp. Bot. 56, 865–877. doi: 10.1093/jxb/eri079

Scheirs, J., De Bruyn, L. (2005). Plant-mediated effects of drought stress on host preference and performance of a grass miner. Oikos 108, 371–385. doi: 10.1111/j.0030-1299.2005.13715.x

Scheirs, J., De Bruyn, L., Verhagen, R. (2001). Nutritional benefits of the leaf-mining behaviour of two grass miners: A test of the selective feeding hypothesis. Ecol. Entomol. 26, 509–516. doi: 10.1046/j.1365-2311.2001.00356.x

Schoonhoven, L.M., Van Loon, J.J.A., Dicke, M. (2005). Insect-plant biology. Oxford university press.

Schwachtje, J., Baldwin, I.T. (2008). Why does herbivore attack reconfigure primary metabolism? Plant Physiol. 146, 845–851. doi: 10.1104/pp.107.112490

Sieciechowicz, K.A., Joy, K.W., Ireland, R.J. (1988). The metabolism of asparagine in plants. Phytochemistry 27, 663–671. doi: 10.1016/0031-9422(88)84071-8

Sømme, L. (1965). Changes in sorbitol content and supercooling points in overwintering eggs of the European red mite (*Panonychus ulmi* (koch)). Can. J. Zool. 43, 881–884. doi: 10.1139/z65-089

Stone, G.N., Schönrogge, K. (2003). The adaptative significance of insect gall morphology. Trends Ecol. Evol. 18, 512–522. doi: 10.1016/S0169-5347(03)00247-7

Suzuki, D.K., Fukushi, Y., Akimoto, S.I. (2009). Do aphid galls provide good nutrients for the aphids? Comparisons of amino acid concentrations in galls among *Tetraneura* species (Aphididae: Eriosomatinae). Arthropod-Plant Interact. 3, 241–247. doi: 10.1007/s11829-009-9064-9

Van Handel, E. (1985). Rapid determination of glycogen and sugars in mosquitoes. J. Am. Mosq. Control Assoc. 1, 299–301.

Van Zyl, L., Ferreira, A.V. (2003). Amino acid requirements of springbok (*Antidorcas marsupialis*), blesbok (*Damaliscus dorcas phillipsi*) and impala (*Aepyceros melampus*) estimated by the whole empty body essential amino acid profile. Small Ruminant Res. 47, 145–153. doi: 10.1016/S0921-4488(02)00248-1

Walters, D.R., McRoberts, N. (2006). Plants and biotrophs: A pivotal role for cytokinins? Trends Plant Sci. 11, 581–586. doi: 10.1016/j.tplants.2006.10.003

Whiteman, N.K., Groen, S.C., Chevasco, D., Bear, A., Beckwith, N., Gregory, T.R., Denoux, C., Mammarella, N., Ausubel, M., Pierce, N.E. (2011). Mining the plant-herbivore interface with a leafmining *Drosophila* of *Arabidopsis*. Mol. Ecol. 20, 995–1014. doi: 10.1111/j.1365-294X.2010.04901.x

Wolfe, G.R., Hendrix, D.L., Salvucci, M.E. (1998). A thermoprotective role for sorbitol in the silverleaf whitefly, *Bemisia argentifolii*. J. Insect Physiol. 44, 597–603. doi: 10.1016/S0022-1910(98)00035-3

Wu, D., Daugherty, S.C., Van Aken, S.E., Pai, G.H., Watkins, K.L., Khouri, H., Tallon, L.J., Zaborsky, J.M., Dunbar, H.E., Tran, P.L., Moran, N.A., Eisen, J.A. (2006). Metabolic complementarity and genomics of the dual bacterial symbiosis of sharpshooters. PloS Biol. 4, e188. doi: 10.1371/journal.pbio.0040188

Wyatt, G.R. (1967). The biochemistry of sugars and polysaccharides in insects. Adv. Insect Physiol. 4, 287–360. doi: 10.1016/S0065-2806(08)60210-6

Zhang, H., Dugé De Bernonville, T., Body, M., Glevarec, G., Reichelt, M., Unsicker, S., Bruneau, M., Renou, J.-P., Huguet, E., Dubreuil, G., Giron, D. (2016). Leaf-mining by *Phyllonorycter blancardella* reprograms the host-leaf transcriptome to modulate phytohormones associated with nutrient mobilization and plant defense. J. Insect Physiol. 84, 114–127. doi: 10.1016/j.jinsphys.2015.06.003

Zhang, H., Guiguet, A., Dubreuil, G., Kisiala, A., Andreas, P., Emery, R.J., Huguet, E., Body, M., Giron, D. (2017). Dynamics and origin of cytokinins involved in plant manipulation by a leaf-mining insect. Insect Sci. 24, 1065–1078. doi: 10.1111/1744-7917.12500

Zhou, S., Lou, Y.R., Tzin, V., Jander, G. (2015). Alteration of plant primary metabolism in response to insect herbivory. Plant Physiol. 169, 1488–1498. doi: 10.1104/pp.15.01405

