## Supplements for "Manipulation of plant primary metabolism by leaf-mining larvae in the race against leaf senescence"

**Supplement 1.** Quantification of eaten tissues by gravimetry and correction for sugar content. The withdrawal of sugar-rich mesophyll tissues by leaf-mining insects, and the over-representation of sugar-free epidermis in the mined-tissue samples, have been taken into account when comparing mined and unmined tissues. Thus, gravimetry was used to estimate the amount of mesophyll eaten by tissue-feeding larvae, which in turn allowed us to correct biochemical data accordingly. Unmined areas similar to mined areas (Figure S1) were dissected in leaves for intermediate and late stage (Figure S2). Following lyophilisation (Bioblock Scientific Alpha1-4LDplus lyophilizator), leaf samples were weighted (Mettler-Toledo micro-balance model ME30, Mettler-Toledo, Viroflay, France). Regression curves were used to correct biochemical data (Figure S3).

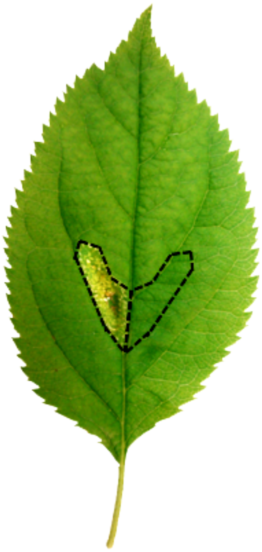

**Figure S1.** Similar areas of mined (left) and unmined (right) tissues dissected in apple-tree leaves.

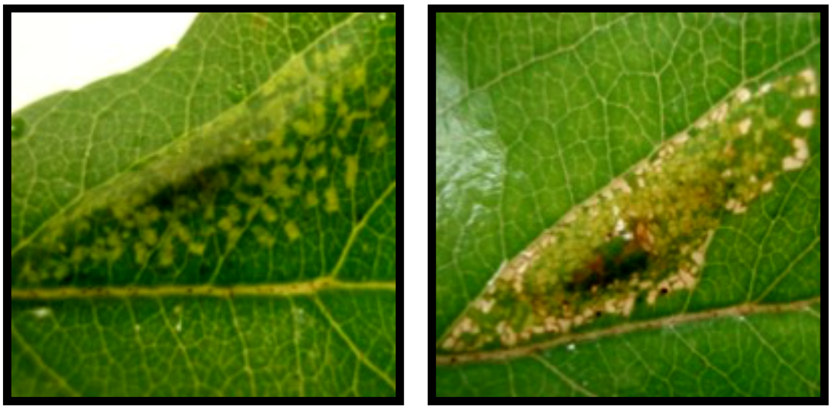

**Figure S2.** Mined tissues. Right: Intermediate stage (left; L4 instar) with spongy parenchyma cells consumed leading to light green patches. Late stage (right; L5 instar) with spongy and palisade parenchyma cells consumed resulting in white and translucent patches where only the epidermis subsists (see Body et al., 2015 for details).

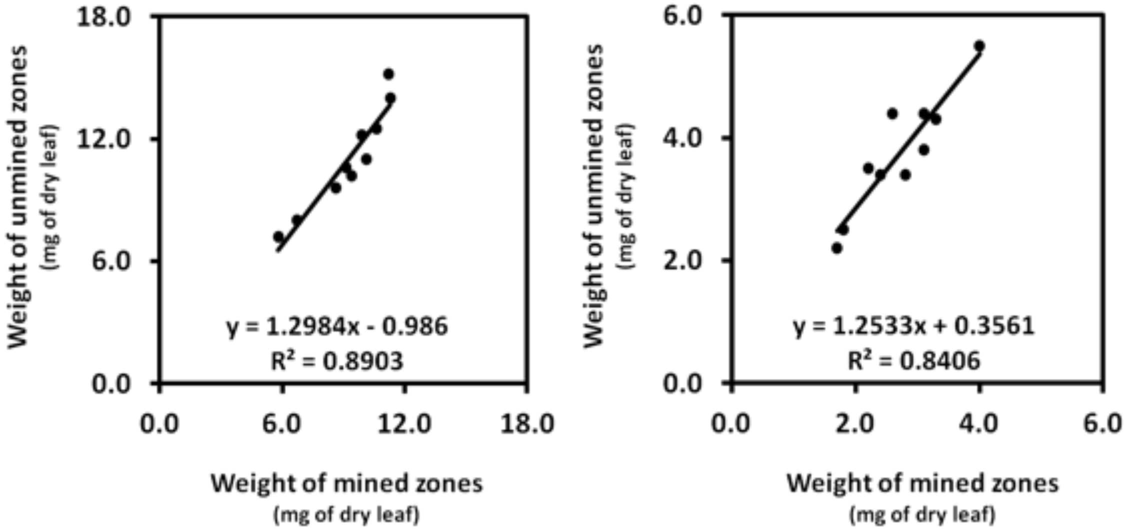


**Figure S3.** Regression curves used to correct data for intermediate (left) and late stage (right).

**Reference**

Body, M., V. Burlat, and D. Giron. 2015. Hypermetamorphosis in a leaf-miner allows insects to cope with a confined nutritional space. Arthropod-Plant Interactions 9:75–84.

**Supplement 2.** Statistical results for sugar contents (starch, total soluble and individual sugars) and sugar composition between unmined and mined tissues for both fluid- and tissue-feeding instars both on green and on yellow leaves. In all cases, *Behrens-Fisher post-hoc tests* were conducted for nutrient quantities and *MANOVA* for sugar compositions (* = $P < 0.05$; ** = $P < 0.01$; *** = $P < 0.001$; ns = no significant difference).

### A. Effect of senescence

| **Green vs. yellow leaves** | **Unmined control tissue** | **Tissue mined by fluid-feeding stage** (early instars) | **Tissue mined by tissue-feeding stage** (late instars) |
|---|---|---|---|
| Starch | P = 0.033 * | P = 0.945 ns | P = 0.001 ** |
| Total soluble sugars | P = 0.000 *** | P = 0.773 ns | P = 0.936 ns |
| Sorbitol | P = 0.000 *** | P = 0.000 *** | P = 0.426 ns |
| Sucrose | P = 0.000 *** | P = 0.024 * | P = 0.999 ns |
| Glucose | P = 0.001 ** | P = 0.888 ns | P = 0.999 ns |
| Fructose | P = 0.027 * | P = 0.001 ** | P = 0.999 ns |
| Sugar composition | $F_{4,75}$ = 18.49, P = 0.000 *** | $F_{4,25}$ = 47.86, P = 0.000 *** | $F_{4,45}$ = 0.99, P = 0.423 ns |

### B. Effect of fluid-feeders

| **Unmined vs. mined tissues** | **Green leaf** | **Yellow leaf** |
|---|---|---|
| Starch | P = 0.000 *** | P = 0.229 ns |
| Total soluble sugars | P = 0.033 * | P = 0.000 *** |
| Sorbitol | P = 1.000 ns | P = 0.009 ** |
| Sucrose | P = 0.732 ns | P = 0.000 *** |
| Glucose | P = 0.807 ns | P = 0.000 *** |
| Fructose | P = 0.000 *** | P = 0.055 ns |
| Sugar composition | $F_{4,50}$ = 5.25, P = 0.001 ** | $F_{4,50}$ = 1.16, P = 0.341 ns |

### C. Effect of tissue-feeders

| **Unmined vs. mined tissues** | **Green leaf** | **Yellow leaf** |
|---|---|---|
| Starch | P = 0.999 ns | P = 0.835 ns |
| Total soluble sugars | P = 0.893 ns | P = 0.000 *** |
| Sorbitol | P = 0.999 ns | P = 0.000 *** |
| Sucrose | P = 0.756 ns | P = 0.000 *** |
| Glucose | P = 0.068 ns | P = 0.460 ns |
| Fructose | P = 0.578 ns | P = 0.873 ns |
| Sugar composition | $F_{4,60}$ = 0.82, P = 0.517 ns | $F_{4,60}$ = 9.92, P = 0.000 *** |

### D. Effect of leaf-miner ontogeny

| **Fluid- vs. tissue-feeders** | **Green leaf** | **Yellow leaf** |
|---|---|---|
| Starch | P = 0.000 *** | P = 0.858 ns |
| Total soluble sugars | P = 0.707 ns | P = 0.534 ns |
| Sorbitol | P = 0.999 ns | P = 0.013 * |
| Sucrose | P = 0.341 ns | P = 0.994 ns |
| Glucose | P = 0.999 ns | P = 0.415 ns |
| Fructose | P = 0.061 ns | P = 0.743 ns |
| Sugar composition | $F_{4,35}$ = 3.31, P = 0.021 * | $F_{4,35}$ = 6.93, P = 0.000 *** |

**Supplement 3.** Statistical results for protein-bound amino acid contents between unmined and mined tissues for both fluid- and tissue-feeding instars both on green and on yellow leaves. In all cases, *Behrens-Fisher post-hoc tests* were conducted (* = $P < 0.05$; ** = $P < 0.01$; *** = $P < 0.001$; ns = no significant difference).

### A. Effect of senescence

| **Green vs. yellow leaves** | **Unmined control tissue** | **Tissue mined by fluid-feeding stage** (early instars) | **Tissue mined by tissue-feeding stage** (late instars) |
|---|---|---|---|
| Ala | P = 0.000 *** | P = 0.999 ns | P = 0.901 ns |
| Gly | P = 0.000 *** | P = 0.999 ns | P = 0.928 ns |
| Val | P = 0.000 *** | P = 0.209 ns | P = 0.999 ns |
| Leu | P = 0.000 *** | P = 0.050 ns | P = 0.992 ns |
| Ile | P = 0.000 *** | P = 0.967 ns | P = 0.698 ns |
| Thr + Ser | P = 0.986 ns | P = 0.994 ns | P = 0.973 ns |
| Pro | P = 0.000 *** | P = 0.999 ns | P = 0.510 ns |
| Asn | P = 0.001 ** | P = 0.935 ns | P = 1.000 ns |
| Arg + Asp | P = 0.999 ns | P = 0.000 *** | P = 0.810 ns |
| Met | P = 0.989 ns | P = 0.203 ns | P = 0.632 ns |
| Phe + Glu | P = 0.000 *** | P = 1.000 ns | P = 0.038 * |
| Cys | P = 0.148 ns | P = 0.789 ns | P = 0.117 ns |
| Gln | P = 0.013 * | P = 0.074 ns | P = 1.000 ns |
| Lys | P = 0.000 *** | P = 0.940 ns | P = 0.000 *** |
| His | P = 0.000 *** | P = 0.651 ns | P = 0.024 * |
| Tyr | P = 0.000 *** | P = 0.908 ns | P = 0.015 * |
| Trp | P = 0.999 ns | P = 0.885 ns | P = 0.477 ns |
| Total | P = 0.000 *** | P = 0.983 ns | P = 0.955 ns |
| EAA | P = 0.000 *** | P = 0.781 ns | P = 0.346 ns |
| NEAA | P = 0.000 *** | P = 0.999 ns | P = 1.000 ns |

### B. Effect of fluid-feeders

| **Unmined vs. mined tissues** | **Green leaf** | **Yellow leaf** |
|---|---|---|
| Ala | P = 0.588 ns | P = 0.129 ns |
| Gly | P = 0.279 ns | P = 0.013 ** |
| Val | P = 0.284 ns | P = 0.027 * |
| Leu | P = 0.904 ns | P = 0.000 *** |
| Ile | P = 0.933 ns | P = 0.000 *** |
| Thr + Ser | P = 0.624 ns | P = 0.000 *** |
| Pro | P = 0.984 ns | P = 0.585 *** |
| Asn | P = 0.203 ns | P = 0.009 ** |
| Arg + Asp | P = 0.840 ns | P = 0.000 *** |
| Met | P = 0.999 ns | P = 0.019 * |
| Phe + Glu | P = 0.541 ns | P = 0.000 *** |
| Cys | P = 0.999 ns | P = 0.616 ns |
| Gln | P = 1.000 ns | P = 0.000 *** |
| Lys | P = 0.051 ns | P = 0.013 * |
| His | P = 0.059 ns | P = 0.000 *** |
| Tyr | P = 0.715 ns | P = 0.000 *** |
| Trp | P = 0.486 ns | P = 0.136 ns |
| Total | P = 0.998 ns | P = 0.000 *** |
| EAA | P = 0.999 ns | P = 0.000 *** |
| NEAA | P = 0.757 ns | P = 0.000 *** |

### C. Effect of tissue-feeders

| Unmined vs. mined tissues | Green leaf | Yellow leaf |
|---|---|---|
| Ala | P = 0.261 ns | P = 0.000 *** |
| Gly | P = 0.378 ns | P = 0.001 ** |
| Val | P = 1.000 ns | P = 0.000 *** |
| Leu | P = 0.836 ns | P = 0.000 *** |
| Ile | P = 0.998 ns | P = 0.000 *** |
| Thr + Ser | P = 0.537 ns | P = 0.000 *** |
| Pro | P = 0.011 * | P = 0.000 *** |
| Asn | P = 0.041 * | P = 0.000 *** |
| Arg + Asp | P = 0.152 ns | P = 0.000 *** |
| Met | P = 0.246 ns | P = 0.000 *** |
| Phe + Glu | P = 0.810 ns | P = 0.000 *** |
| Cys | P = 0.984 ns | P = 0.027 * |
| Gln | P = 0.515 ns | P = 0.003 ** |
| Lys | P = 0.999 ns | P = 0.001 ** |
| His | P = 0.999 ns | P = 0.000 *** |
| Tyr | P = 0.992 ns | P = 0.000 *** |
| Trp | P = 0.174 ns | P = 0.015 * |
| Total | P = 0.526 ns | P = 0.000 *** |
| EAA | P = 0.996 ns | P = 0.000 *** |
| NEAA | P = 0.128 ns | P = 0.000 *** |

### D. Effect of leaf-miner ontogeny

| Fluid- vs. tissue-feeders | Green leaf | Yellow leaf |
|---|---|---|
| Ala | P = 1.000 ns | P = 1.000 ns |
| Gly | P = 0.999 ns | P = 0.995 ns |
| Val | P = 0.469 ns | P = 0.092 ns |
| Leu | P = 0.992 ns | P = 0.926 ns |
| Ile | P = 0.889 ns | P = 0.895 ns |
| Thr + Ser | P = 0.995 ns | P = 0.776 ns |
| Pro | P = 0.321 ns | P = 0.157 ns |
| Asn | P = 0.997 ns | P = 1.000 ns |
| Arg + Asp | P = 0.128 ns | P = 0.837 ns |
| Met | P = 0.674 ns | P = 0.999 ns |
| Phe + Glu | P = 0.392 ns | P = 0.999 ns |
| Cys | P = 1.000 ns | P = 0.999 ns |
| Gln | P = 0.921 ns | P = 0.004 ** |
| Lys | P = 0.050 ns | P = 0.999 ns |
| His | P = 0.062 ns | P = 0.910 ns |
| Tyr | P = 0.485 ns | P = 0.997 ns |
| Trp | P = 0.999 ns | P = 0.999 ns |
| Total | P = 0.999 ns | P = 1.000 ns |
| EAA | P = 0.995 ns | P = 0.997 ns |
| NEAA | P = 1.000 ns | P = 1.000 ns |

**Supplement 4.** Statistical results for free amino acid contents between unmined and mined tissues for both fluid- and tissue-feeding instars both on green and on yellow leaves. In all cases, *Behrens-Fisher post-hoc tests* were conducted (* = $P$ < 0.05; ** = $P$ < 0.01; *** = $P$ < 0.001; ns = no significant difference).

**A. Effect of senescence**

| **Green vs. yellow leaves** | **Unmined control tissue** | **Tissue mined by fluid-feeding stage** (early instars) | **Tissue mined by tissue-feeding stage** (late instars) |
|---|---|---|---|
| Ala | P = 0.000 *** | P = 0.000 *** | P = 1.000 ns |
| Gly | P = 0.000 *** | P = 0.000 *** | P = 0.863 ns |
| Val | P = 0.000 *** | P = 0.000 *** | P = 0.000 *** |
| Leu | P = 0.000 *** | P = 0.000 *** | P = 0.999 ns |
| Ile | P = 0.000 *** | P = 0.000 *** | P = 0.000 *** |
| Thr + Ser | P = 0.183 ns | P = 0.000 *** | P = 0.408 ns |
| Pro | P = 0.000 *** | P = 0.000 *** | P = 0.999 ns |
| Asn | P = 0.000 *** | P = 0.000 *** | P = 0.727 ns |
| Arg + Asp | P = 0.010 * | P = 0.000 *** | P = 0.582 ns |
| Met | --- | --- | --- |
| Phe + Glu | P = 0.007 ** | P = 0.001 ** | P = 0.909 ns |
| Cys | --- | --- | --- |
| Gln | P = 0.000 *** | P = 0.001 ** | P = 0.954 ns |
| Lys | P = 0.999 ns | P = 0.002 ** | P = 0.998 ns |
| His | --- | --- | --- |
| Tyr | P = 0.013 * | P = 0.997 ns | P = 0.490 ns |
| Trp | P = 0.000 *** | P = 0.000 *** | P = 0.001 ** |
| Total | P = 0.000 *** | P = 0.000 *** | P = 0.837 ns |
| EAA | P = 0.000 *** | P = 0.000 *** | P = 0.064 ns |
| NEAA | P = 0.000 *** | P = 0.000 *** | P = 0.884 ns |

**B. Effect of fluid-feeders**

| **Unmined vs. mined tissues** | **Green leaf** | **Yellow leaf** |
|---|---|---|
| Ala | P = 0.000 *** | P = 0.627 ns |
| Gly | P = 0.000 *** | P = 0.966 ns |
| Val | P = 0.000 *** | P = 0.547 ns |
| Leu | P = 0.000 *** | P = 0.999 ns |
| Ile | P = 0.000 *** | P = 0.768 ns |
| Thr + Ser | P = 0.999 ns | P = 0.202 ns |
| Pro | P = 0.000 *** | P = 0.961 ns |
| Asn | P = 0.937 ns | P = 0.000 *** |
| Arg + Asp | P = 0.000 *** | P = 0.000 *** |
| Met | --- | --- |
| Phe + Glu | P = 0.000 *** | P = 0.194 ns |
| Cys | --- | --- |
| Gln | P = 0.784 ns | P = 0.002 ** |
| Lys | P = 0.954 ns | P = 0.001 ** |
| His | --- | --- |
| Tyr | P = 1.000 ns | P = 0.728 ns |
| Trp | P = 0.085 ns | P = 0.192 ns |
| Total | P = 0.505 ns | P = 0.000 *** |
| EAA | P = 0.000 *** | P = 0.000 *** |
| NEAA | P = 0.879 ns | P = 0.000 *** |

**C. Effect of tissue-feeders**

| Unmined vs. mined tissues | Green leaf | Yellow leaf |
|---|---|---|
| Ala | P = 0.999 ns | P = 0.026 * |
| Gly | P = 0.009 ** | P = 0.620 ns |
| Val | P = 0.020 * | P = 0.062 ns |
| Leu | P = 0.160 ns | P = 0.004 ** |
| Ile | P = 0.003 ** | P = 0.008 ** |
| Thr + Ser | P = 0.604 ns | P = 0.974 ns |
| Pro | P = 0.685 ns | P = 0.056 ns |
| Asn | P = 0.000 *** | P = 0.359 ns |
| Arg + Asp | P = 0.041 * | P = 0.007 ** |
| Met | --- | --- |
| Phe + Glu | P = 0.267 ns | P = 0.999 ns |
| Cys | --- | --- |
| Gln | P = 0.466 ns | P = 0.093 ns |
| Lys | P = 1.000 ns | P = 0.999 ns |
| His | --- | --- |
| Tyr | P = 0.979 ns | P = 0.871 ns |
| Trp | P = 0.351 ns | P = 0.289 ns |
| Total | P = 0.001 ** | P = 0.174 ns |
| EAA | P = 0.045 * | P = 0.996 ns |
| NEAA | P = 0.002 ** | P = 0.115 n |

### D. Effect of leaf-miner ontogeny

| Fluid- vs. tissue-feeders | Green leaf | Yellow leaf |
|---|---|---|
| Ala | P = 0.000 *** | P = 0.994 ns |
| Gly | P = 0.000 *** | P = 0.999 ns |
| Val | P = 0.170 ns | P = 0.003 ** |
| Leu | P = 0.000 *** | P = 0.001 ** |
| Ile | P = 0.999 ns | P = 0.004 ** |
| Thr + Ser | P = 0.739 ns | P = 0.116 ns |
| Pro | P = 0.000 *** | P = 0.056 ns |
| Asn | P = 0.002 ** | P = 0.173 ns |
| Arg + Asp | P = 0.000 *** | P = 0.039 * |
| Met | --- | --- |
| Phe + Glu | P = 0.002 ** | P = 0.472 ns |
| Cys | --- | --- |
| Gln | P = 0.607 ns | P = 0.063 ns |
| Lys | P = 0.999 ns | P = 0.008 ** |
| His | --- | --- |
| Tyr | P = 0.986 ns | P = 0.698 ns |
| Trp | P = 0.240 ns | P = 0.999 ns |
| Total | P = 0.000 *** | P = 0.009 ** |
| EAA | P = 0.000 *** | P = 0.020 * |
| NEAA | P = 0.003 ** | P = 0.034 * |
